# CD4+ T Cell Depletion Before Primary Dengue and/or Secondary Zika Infection Reveals Mechanistic Correlates of Antibody Functionality in Rhesus Macaques

**DOI:** 10.1101/2024.02.22.580962

**Authors:** Crisanta Serrano-Collazo, Angel Miranda, Lorna A. Cruz, Sandra Henein, Mitchell Sanchez-Rosado, Laura Alvarez, Teresa Arana, Melween I. Martinez, Chiara Roman, Armando G. Burgos, Aravinda de Silva, Carlos A. Sariol

## Abstract

Dengue (DENV) virus and Zika virus (ZIKV) are two flaviviruses of major public health concern. One drawback designing effective vaccines is our limited understanding of the mechanisms ruling protection or harm among DENV serotypes, or between DENV and ZIKV. Here, we depleted rhesus macaques of CD4^+^ T cells *in vivo* before primary DENV infection and/or secondary ZIKV challenge to recreate a sub-optimal priming of the humoral immune response. Our results support that CD4^+^ T cells are needed to induce a quantitative and type-specific effective humoral immune response against primary DENV, but also against secondary ZIKV in DENV-experimented subjects. Our results also indicate a limited contribution of the DENV-Memory B cells to anti-ZIKV response. Furthermore, our results suggest that a suboptimal B cell priming during a primary DENV infection does differentially impact different antibody (Abs) properties. While binding or neutralization of ZIKV or DENV during a subsequent exposure to ZIKV is not affected by the lack of CD4^+^ T - B cells interaction during a primary DENV infection, that interaction is critical to guarantee the Abs specificity. Also, we found that depleting CD4^+^ T cells before DENV primary infection but not before ZIKV challenge significantly increases Abs cross-reactivity against DENV-EDIII domain and DENV-NS1 protein but not against ZIKV-EDIII domain or NS1 protein. Furthermore, there was more cross-reactivity among the DENV-NS1 proteins than against DENV-EDIII domains, suggesting that during a primary DENV infection CD4^+^ T cells have a different weight in the responses against EDIII domain and NS1 protein. The proper Abs binding and neutralization with increased cross-reactivity profile was associated with limited frequency of circulating peripheral T helper cells (pTfh) with T helper 1 phenotype (CD4+/CXCR5+/CXCR3+) and expressing markers related to B cell activation (CXCR5+/CXCR3+/PD-1+/ICOS+) in the group depleted of CD4^+^ T cells only before primary DENV infection. However, memory B cells – but not Antibody Secreting Cells (ASC) activation 7 days after the infection – positively correlate with those two populations of pTfh. Finally, when Abs cross-reactivity values were incorporated in a Principal Component Analysis (PCA), the DENV-CD4^+^ T depleted group separates from the other two groups with similar Abs binding and neutralization profiles. Our result strongly suggests that during a heterologous sequential DENV/ZIKV infections Abs binding, and neutralization, may be regulated by different factors than their specificity. Before, the induction of cross-neutralizing Abs has been described in the context of secondary DENV infection. Here, for the first time, we are reproducing the experimental conditions leading to the generation of such Abs population *in vivo*. In summary, we show that suboptimal immune priming during a primary flavivirus infection has functional consequences during a secondary heterologous infection. Finally, we shown that CD8^+^ T cells are essential to guarantee an optimal Abs neutralization activity. These results have huge implications understanding the immune response to DENV vaccines (and maybe ZIKV), including why an optimal vaccine or natural-induced neutralizing response not necessarily protects or enhances pathogenesis during a subsequent natural heterologous exposure.

## Introduction

Dengue (DENV) and Zika (ZIKV) viruses are flaviviruses of significant public health concern (Li et al., 2021; Paixao et al., 2022; Qiao et al., 2021; Tian et al., 2022; Wong et al., 2022). DENV is responsible for about 390-500 million infections per year, of which 96 million present different levels of disease severity (Bhatt et al., 2013) including Dengue Hemorrhagic Fever/ Shock Syndrome with up to 20 % mortality rate if untreated (WHO/TDR, 2009). It is estimated that the population at risk will increase in 60% by 2080 for a total of 6.1 billion people (Messina et al., 2019). ZIKV infections are usually mild but have been associated with unique severe adverse outcomes such as fetal loss (Campos et al., 2016), congenital ZIKV syndrome (CZS) (Brasil et al., 2016), Guillain-Barré syndrome (GBS) (Cao-Lomeau et al., 2016), and rare cases of encephalopathy (Roze et al., 2016), meningoencephalitis (Carteaux et al., 2016), myelitis (Mecharles et al., 2016), uveitis (Furtado et al., 2016), and severe thrombocytopenia (Sharp et al., 2016).

DENV and ZIKV circulate in similar geographic areas (Braga et al., 2017; Rodriguez-Morales et al., 2016), and the effect of prior DENV or ZIKV immunity to each other is subject of an ongoing debate. Some evidence from animal models and humas suggest that prior DENV immunity has a protective role in secondary ZIKV infection (Ana Carolina Bernardes Terzian, 2017; Breitbach et al., 2019; McCracken et al., 2017; Pantoja et al., 2017). Other works propose that prior DENV infection may increase ZIKV disease severity (Bardina et al., 2017; Crooks et al., 2021; Katzelnick et al., 2020b; Katzelnick et al., 2021). Intermediate pre-existing levels of naturally acquired anti-DENV or anti-ZIKV antibodies (Abs) were found to be associated with enhancement of symptomatic and severe DENV. However, subjects with high anti-DENV or anti-ZIKV Abs are at much lower risk of symptomatic and severe DENV2, with rates similar to DENV-naïve individuals (Katzelnick et al., 2020a). To add complexity to the field, recently it has been documented that different DENV and ZIKV interactions can be different or asymmetrical depending on the DENV serotype {Bos, 2023 #2166;Katzelnick, 2020 #2029}. Nevertheless, the mechanisms supporting a proportionate immune response among all four DENV serotypes or DENV and ZIKV after a natural infection – particularly after vaccination – still remains as a critical knowledge gap.

Immunity to one DENV serotype is believed to confer type specific (TS) protective Abs or immunity to that infecting serotype, but can also induce cross-reacting (CR) Abs able to amplify the pathogenesis during a secondary infection with a different serotype – a phenomenon known as antibody-dependent enhancement (ADE) (Guzman et al., 2013; Halstead, 2003; Halstead et al., 1983; Katzelnick et al., 2017). As a result, DENV vaccines are designed as tetravalent formulations to induce balanced Abs (Abs) response against all four DENV serotypes at the same time. Two of the leading DENV vaccines (Dengvaxia and TAK-003), based on attenuated chimeric viruses (Deauvieau et al., 2007; Guy et al., 2017; Huang et al., 2013; Osorio et al., 2011b) has shown to induce unbalanced immune responses (Biswal et al., 2020; Guirakhoo et al., 2002; Hadinegoro et al., 2015; Halstead, 2018; Sabchareon et al., 2012; White et al., 2021). A third vaccine candidate developed by the Laboratory of Infectious Diseases at the National Institutes of Health (TV003) based on live attenuated DENV viruses (Durbin et al., 2013; Durbin et al., 2011; Whitehead, 2016), induced TS Abs to all four DENV serotypes and protection against challenge (Kirkpatrick et al., 2016; Nivarthi et al., 2021). Recent results from phase 3 are encouraging, but differences in efficacy were still reported between DENV1 [89.5% (95% CI, 78.7 to 95.0)] and DENV2 [69.6% (95% CI, 50.8 to 81.5)] the two serotypes detected in the three years of follow up period (Kallás et al., 2024). The limitations of Dengvaxia and TAK-003 formulations in inducing a balanced immune response to all four DENV serotypes have been associated to unequal replication of vaccine components or to the lack of/incomplete cellular immune response to the nonstructural (NS) proteins, among others (Grifoni et al., 2020; Guy et al., 2008; Guy et al., 2020; Thomas et al., 2009; Waickman et al., 2019a; Waickman et al., 2019b). For Dengvaxia, the DENV4 component was replicating and immunodominant, compared to the other three serotypes (Guy et al., 2009); a similar situation is documented with DENV2 in the TAK-003 vaccine in non-human primates (NHP) and humans (Osorio et al., 2011a; Rupp et al., 2015; Tricou et al., 2020).

Total Abs to each DENV serotypes have been used as a marker of protection for DENV vaccines, as it has shown to correlate with the efficacy of other flavivirus vaccines like yellow fever vaccine (YFV), Japanese encephalitis vaccine (JEV) (Appaiahgari et al., 2006; Goncalvez et al., 2008; Gupta et al., 2008; Raengsakulrach et al., 1999; Saini and Vrati, 2003), and tick-borne encephalitis (TBEV) (Beran et al., 2019; Leonova and Pavlenko, 2009; Maikova et al., 2016; Venturi et al., 2009; Vorovitch et al., 2019). But recent growing evidences point out that TS neutralizing Abs (nAbs) above a certain threshold may be a better indicator of vaccine efficacy than the total Abs (Henein et al., 2021; Nivarthi et al., 2021). Earlier works also provide explanations as of why some Abs might neutralize DENVs in cell culture, but not protect *in vivo* (Raut et al., 2019). Martinez et al. assembled a comprenhensive review on the multiple factors modifiying the efficacy of the immune response to DENV and DENV vaccines, incluidng the intra and inter serotypic viral antigenic diversity and the genotypic diversity among others {Martinez, 2021 #2068}.

All those observations and findings from natural infections and vaccines are supported on Abs response mainly against the structural proteins, reinforcing the concept that immunogenicity of the structural proteins does not guarantee protection from natural infection or vaccination efficacy. Most humoral immunity epitopes are located within the domains of the DENV envelope (E) protein, while cellular immunity epitopes are located on the NS proteins (Pinheiro et al., 2021; Rothman, 2010). Additionally, the NS1 protein is particularly interesting and unique, as it is a protein produced and secreted during viral replication, which make it highly immunogenic (Zhang et al., 2023) inducing significant levels of anti-NS1 Abs in DENV- and ZIKV-infected individuals (Aquino et al., 2023; Castro-Trujillo et al., 2023; Churdboonchart et al., 1991; Costa et al., 2007; Petphong et al., 2023). It is documented that DENV E and NS1 proteins equally score 4 – out of the 10 proteins – in an immunogenicity score (Weiskopf et al., 2016), and some reports have pointed NS1 as a target antigen for the development of DENV vaccines (Costa et al., 2007; Costa et al., 2006; Henchal et al., 1988; Wan et al., 2014), while others suggest a role for this protein in the pathogenesis (Perera et al., 2024; Shu et al., 2000; Zhang et al., 2023). Thus, it still remains a puzzle due to its ability to elicit both protective and potentially pathogenic immune responses. On this work, in addition to the conventional Abs to the structural proteins, we explore the contribution of NS1 to the immune response during the primary DENV infection and a secondary ZIKV challenge. Beyond the cannonical humoral component, a robust T cell response is needed to confer protection after a natural DENV or ZIKV infection (Elong Ngono and Shresta, 2019; Roth et al., 2018; Tian et al., 2019; Weiskopf and Sette, 2014; Yauch et al., 2009) and most of the cognates epitopes for the T cells (CD4^+^ and CD8^+^ T cells) are located in the NS proteins (Grifoni et al., 2017a; Rivino et al., 2013; Rivino and Lim, 2017). Despite this, one common aspect shared among all three leading DENV vaccines is that they do not present NS proteins of all DENV serotypes to the immune system. Dengvaxia is based on a Yellow Fever (YF) 17D backbone and contains no DENV NS proteins (Guirakhoo et al., 2001). The Takeda vaccine uses a DENV-2 backbone and has only DENV-2 NS proteins (Huang et al., 2013; Kinney et al., 1997). The NIH/Butantan/MSD vaccine has full genome DENV-1, -3, and -4 components with a DENV-2/-4 chimera based on a DENV-4 backbone. This vaccine possesses NS proteins from DENV-1, -3, and -4 but not of DENV2 (Blaney et al., 2006; Blaney et al., 2008; Whitehead et al., 2003a; Whitehead et al., 2003b). Further, data on the contribution of T cells and their cognate epitopes on the NS proteins to the vaccine efficacy, and potentially protection, is still restricted (Angelo et al., 2017; Guy et al., 2008; Tricou et al., 2022; Waickman et al., 2019a; Weiskopf et al., 2015a). These data underscore how the contribution of NS proteins and T cells has been understudied in the context of DENV vaccination. Pinheiro et al. completed a comprehensive analysis of B and T cells epitopes shared by all three leading DENV vaccines, and concluded that investing in vaccines that contain most of the epitopes involved in protective immunity (cellular and humoral arms) is an important issue to be considered (Pinheiro et al., 2021). In fact, a few years ago the World Health Organization (WHO) recommended that the cell mediated immunity (CMI) induced by DENV vaccines should be studied in detail (Thomas et al., 2009).

Also, it has been shown that CD4^+^ T cells are essential for inducing a protective immune response against DENV and ZIKV (Elong Ngono et al., 2019; Grifoni et al., 2017b; Grifoni et al., 2017c; Hassert et al., 2018; Manh et al., 2020; Weiskopf et al., 2015b; Wen et al., 2020); these cells play a critical role connecting the cellular and humoral immune responses though unique interactions with B cells (Crotty, 2015; Swain et al., 2012). Moreover, immunological characterization of T cells has identified the presence of follicular CD4^+^ T helper cells (Tfh) or “Tfh-like” in blood, known as circulating (cTfh) or peripheral (pTfh) cells. These cells have been described, in terms of function, as the equivalent counterpart of germinal center (GC) Tfh cells in peripheral blood (Bentebibel et al., 2013; Crotty, 2014; Morita et al., 2011). However, there is limited literature available on the role of pTfh cells in DENV (Haltaufderhyde et al., 2018; Izmirly et al., 2022; Wijesinghe et al., 2020) and ZIKV vaccination or infections (Liang et al., 2019; Pardy et al., 2022). Despite their critical role helping B cells form an optimal humoral immune response, there are scarce works studying the direct interactions of CD4^+^ T cells/pTfh cells subsets with B cells and NS proteins, and its effect in Abs functionality against DENV and/or ZIKV (Elong Ngono et al., 2019; Rivino et al., 2013; Rouers et al., 2021). Using a NHP model – rhesus macaques – we asked how the lack of essential cells, such as CD4^+^ T cells, impacts the quality of the humoral immune response (including the response to NS1 protein) during a primary DENV2 infection and its consequences in the response to a secondary heterologous ZIKV infection. We believe this is the type of mechanistic questions that needs to be addressed in order to identify correlates of protections and ultimately design more efficient flavivirus vaccines.

In this work, we depleted rhesus macaques of CD4^+^ T cells before a primary DENV2 infection and/or before a subsequent ZIKV infection one year apart. A cohort was depleted of CD8^+^ T cells before the secondary challenge with ZIKV.

We found that the lack of CD4^+^ T cells before infections results in a significant increase in DENV viremia and DENV-NS1 antigenemia and ZIKV viremia. Lack of CD4^+^ T cells before DENV or ZIKV also results in a delay in total Abs switch documented by limited IgM and IgG production and in a significantly lower anti-DENV and anti-ZIKV NS1 Abs response. We also found that Abs binding or neutralizing DENV or ZIKV after heterologous challenge with ZIKV secondary infection were minimally impacted by the lack of CD4^+^ T cells during DENV primary infection. Nevertheless, the presence of these cells during the primary DENV infection were critical to preserve the specificity of the Abs against the DENV, but not ZIKV EDIII domain and NS1 protein after ZIKV challenge.

This profile of Abs functionality (proper binding and neutralization with increased cross-reactivity) was associated with a significant lower frequency of pTfh TH1 and expression of markers related to B cell activation – PD-I and ICOS – when CD4^+^ T cells were not present during the primary DENV infection. Furthermore, the expression of these markers correlates with the activation of the MBC, but not ASC, in the same group. Interestingly, the depletion of CD8^+^ T cells abrogates the direct correlation between those CD4^+^ T cells subsets and MBC or ASC activation.

The regulation of the antibody response and its quality by CD8^+^ T cells is significantly less characterized than by CD4^+^ T cells. While CD8^+^ T cells are well defined for their cytotoxic response, recent findings have identified a population of CD8^+^ T cells that show distinct phenotypical characteristics of Tfh cells and can be found in the B cell follicles (Chen et al., 2023). These cell subsets show high expression of markers such as CXCR5 (related to follicular-homing) and CXCR3 while downregulating T zone homing markers such as CCR7 (Quigley et al., 2007). Here we show that depleting CD8^+^ T cells eliminates the strong correlation between CD4^+^ T Cells and Abs neutralization activity.

With our experimental design, we could reproduce *in vivo*, first the time, the mechanism inducing those CR nAbs. This Abs population was described in the context of secondary DENV infections (Patel et al., 2017) and has been at the center of an active debate on their protective or pathogenic contribution to the flavivirus response. Our findings are critical to understand the efficacy limitations of Abs induced by current approved DENV vaccines or protecting from natural infections and thus are essential to guide the new generations of DENV and ZIKV vaccines. We also provide an animal model to reproduce an immunological signature of the response to flaviviruses that can be used to test and challenge the protective or disease-enhancing attributes of current and new vaccine candidates or, as in this work, to better characterize flavivirus interactions.

We are unaware of any prior work addressing the complex interactions among CD4^+^ T cells, B cells and NS proteins in response to flavivirus infection in NHPs or correlating the CD8^+^ T cells with the quality of the humoral immune response to flavivirus.

## Results

### Rhesus macaque cohorts, T cell depletion and sample collection

Five groups of flavivirus-naïve rhesus macaques (*Macaca mulatta*) (SFig 1). Prior to any experimental activity, all animals were put through a quarantine period of 40 days. Depletion of T cells was performed by initial subcutaneous (s.c.) administration of 50 mg/kg of anti-CD4+ treatment at 15 days pre-challenge, followed by two intravenous (i.v.) administrations of 7.5 mg/kg at 12- and 9-days pre-challenge. Animals were depleted of CD4+ or CD8 T+ cells, or administered phosphate buffer saline (PBS), followed by a primary infection with DENV-2 (5×10^5^ pfu s.c.) and a subsequent ZIKV challenge (1 × 10^6^ pfu s.c.) one year later, after a second round of T cell depletion – or administration of PBS – before infection. CD4+ T cell depletion was performed on groups 1 and 3 prior to primary DENV2 infection and on groups 1 and 2 prior to secondary ZIKV infection (group 1: n=6, denoted CD4-/DV2/CD4-/ZV; group 2: n=6, denoted CD4+/DV2/CD4-/ZV; group 3: n=6, denoted CD4-/DV2/CD4+/ZV). Group 4 is CD4-competent but CD8-depleted prior to secondary ZIKV infection (n=6, denoted CD4+/DV2/CD8-/ZV). Group 5, the control group, was treated with PBS before each infection (n=5, denoted CD4+/DV2/CD4+/ZV).

### Depletion of CD4+ T cells elicits a CD8+ T cell overcompensation, and vice-versa

To determine if the depletion treatment was effective, frequency of CD4+ and CD8+ T cells was assessed by immunophenotyping using flow cytometry during pre-treatment (pre-TX), post-treatment (post-TX), and post-primary DENV2 infection. Before depletion, CD4+ T cell frequency was similar in all animals and decreased by day 2 post-TX, resulting in a depletion of 99.8% (SFig 2A). This reduction was consistent until day 7 post infection (p.i.), when they began to slowly recover by day 15 post infection p.i.. Further, the absence of CD4+ T cells in depleted animals promoted an increase in CD8+ T cell frequency (SFig 2B).

CD4+ and CD8+ T cell frequencies were also measured before, during and after depletions prior to and during ZIKV challenge (SFig 2C and D). Similar to the prior scenario, CD4+ T cell frequency was similar in all animals, and decreased by day 2 after treatment start (SFig 2C). Interestingly, animals that underwent CD8+ T cell depletion experimented an increase in CD4+ T cell frequency (SFig 2C). Lastly, CD8+ T cells behaved similarly to CD4+ T cells after depletion, with frequency decreasing by day 2 post-TX and beginning to recover by day 7 post-challenge (p.c.), while CD4-depleted animals showed an increase in CD8+ T cell frequency (SFig 2D).

### CD4+ T cell depletion impairs DENV2 viral clearance in flavivirus-naïve macaques

To determine if CD4+ T cell depletion enhances or reduces DENV2 replication, and how it changes depending on various depletion conditions, DENV2 RNAemia was measured in serum during the first 15 days and day 30 after primary DENV2 infection. RNAemia was defined as follows: early from days 1 to 3 p.i., mid from days 4 to 7 p.i., and late from day 7 p.i. onwards. During the early period, viral RNA detection increased similarly in all groups (Fig 1A). By day 5 p.i., the control group showed a decrease in viremia leading to resolution, while the CD4-depleted group still had high detectable RNAemia levels. This difference was statistically significant on days 10 and 11 p.i. (p= 0.000078 and p= 0.001346, respectively). By day 12 p.i. there was no viral RNA detection in any animal from the control group, indicating a faster RNAemia resolution in this group compared to CD4-depleted animals. This suggests a delay in viral clearance in animals that are CD4-deficient. By day 15 p.i. all animals tested negative for DENV2 (Fig 1A and Table S1).

**Fig 1.**
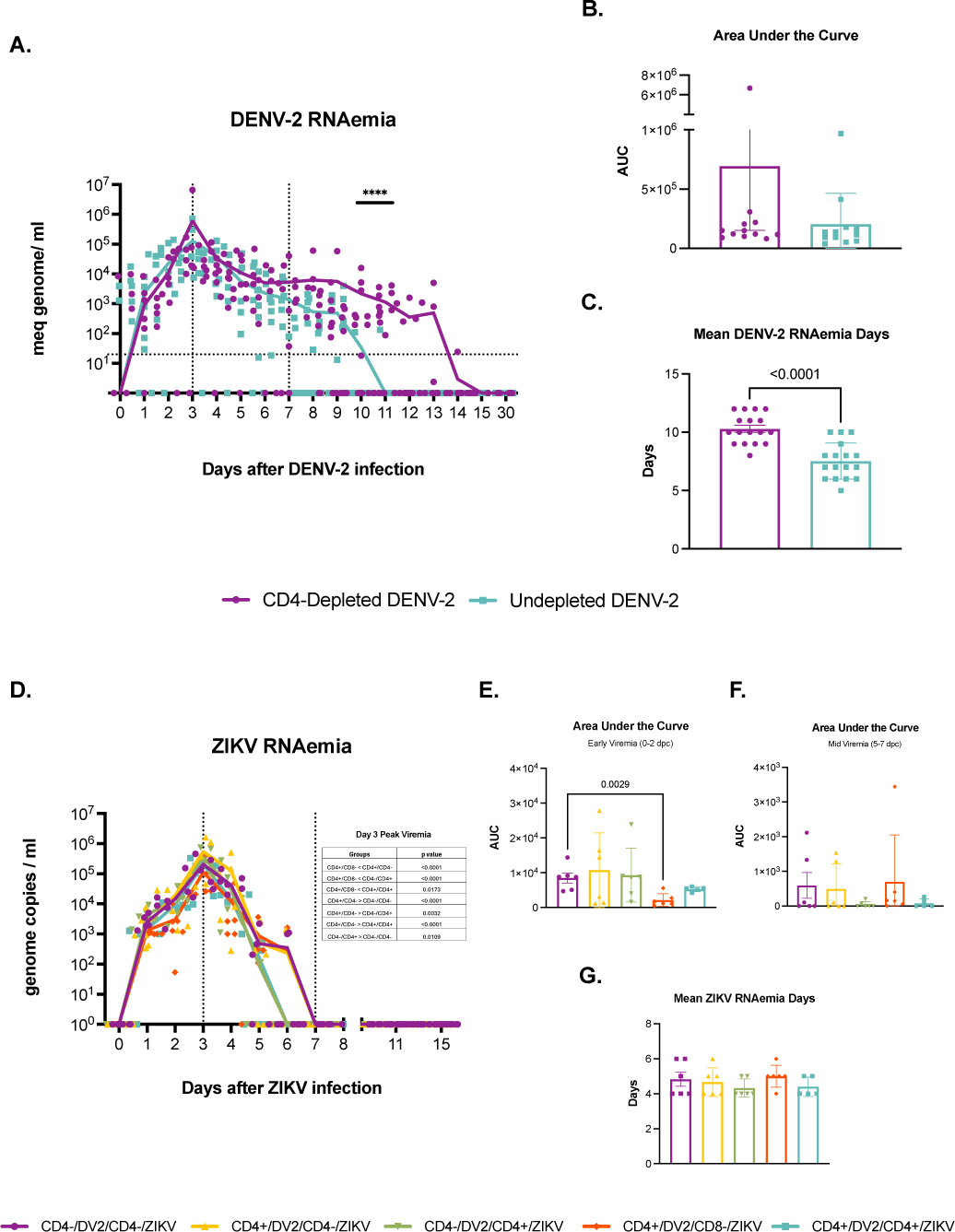
DENV2 and ZIKV RNA kinetics in flavivirus-naïve CD4-depleted and undepleted animals. RNAemia is affected by depletion of CD4+ or CD8+ T cells. **Upper panel**. CD4-depleted animals are depicted in purple and undepleted animals are depicted in turquoise. **(A)** DENV-2 genome copies/mL were measured in serum to monitor viral replication during the first 15 days and day 30 after infection. Genome copies per mL are shown logarithmically. DENV RNAemia was defined as early RNAemia (days 0 to 3 p.i.), mid RNAemia (days 4 to 7 p.i.) and late RNAemia (day 7 p.i. onwards). **(B)** The area under the curve (AUC) was calculated for individual values. **(C)** Average RNAemia days were calculated using the following formula: total viremia days divided by the total number of days during which viremia was monitored. **Lower panel**. Animal cohorts are depicted as follows based on treatment regime: CD4+/DV2/CD4+/ZV (shown in turquoise), CD4-/DV2/CD4+/ZV (shown in green), CD4+/DV2/CD4-/ZV (shown in yellow), CD4-/DV2/CD4-/ZV (shown in purple), and CD4+/DV2/CD8-/ZV (shown in red). **(D)** ZIKV genome copies/mL were measured in serum to monitor viral replication during the first 15 days and day 30 after infection. Genome copies per mL are shown logarithmically. ZIKV RNAemia was defined as early RNAemia (days 0 to 3 p.i.), mid RNAemia (days 4 to 7 p.i.) and late RNAemia (day 7 p.i. onwards). **(E)** The area under the curve (AUC) was calculated for individual values on days 0, 1 and 2 post ZIKV challenge, denominated early viremia. **(F)** The area under the curve (AUC) was calculated for individual values on days 5, 6 and 7 post ZIKV challenge, denominated mid viremia **(G)** Average RNAemia days were calculated using the following formula: total viremia days divided by the total number of days during which viremia was monitored. Statistically significant differences among and within groups were calculated by two-way ANOVA using Tukey’s multiple comparisons test and unpaired multiple t-tests followed by Bonferroni’s multiple comparisons test.

When evaluating the area under the curve (AUC), we found that the CD4-competent animals, although not statistically significant, had lower values, on average, than the CD4-depleted group (Fig 1B). Next, we evaluated the average RNAemia days, defined as the days with detectable viremia divided by the number of animals per group (Fig 1C). The undepleted animals had significantly less mean RNAemia days compared to the depleted animals (p< 0.0001). Taken together, these results suggest that CD4^+^ T cells have an early role controlling DENV2 RNAemia, but it remains unknown if this is due to a direct action or to an interaction with B cells.

### CD4+ but not CD8+ T cells are necessary to control early ZIKV RNAemia in DENV-immune macaques

To assess the impact of T cell depletions in ZIKV replication in DENV2-immune animals, and how it changes depending on various depletion conditions, ZIKV RNAemia was measured in serum during the first 15 days and on day 30 after secondary ZIKV challenge using qRT-PCR. ZIKV RNAemia was defined as follows: early from days 1 to 3 p.i., mid from days 4 to 7 p.i., and late from day 7 p.i. onwards. Although all groups had a similar increase in early viral RNA detection, numerous significant differences were noted on day 3 p.c. which is also when peak viremia was reached (Fig. 1D). The CD4+/DV2/CD8-/ZV group had the lowest peak viremia among all groups, and it was significantly lower in comparison to the CD4+/DV2/CD4-/ZV, CD4-/DV2/CD4+/ZV and CD4+/DV2/CD4+/ZV groups (p< 0.0001, p< 0.0001 and p= 0.0173, respectively). On the other hand, the CD4+/DV2/CD4-/ZV group had a significantly higher peak viremia compared to groups CD4-/DV2/CD4-/ZV, CD4-/DV2/CD4+/ZV and the control group (p< 0.0001, p= 0.0032 and p< 0.0001, respectively). Lastly, the CD4-/DV2/CD4+/ZV had a significantly higher peak viremia compared to the double-depleted group (p= 0.0109) (Fig 1D and Table S2). Of note, historical data from our laboratory shows that ZIKV replication in naive undepleted animals is higher than ZIKV replication in DENV-immune animals regardless of CD4+ T cell status, as well as an earlier peak viremia, which was displaced by one day in the presence of previous DENV2 immunity in the current presented data (Fig. 1D). By day 7 p.c., there was no viral RNA detection in any group.

We observed a difference when comparing all five groups based on AUC on different RNAemia periods (Figures 1E and F). During early RNAemia, CD8-depleted animals had lower AUC levels compared to all groups, and they were significantly lower than the double-depleted animals (p= 0.0029) (Fig 1 E). However, a shift occurs during mid RNAemia where CD8-depleted animals, although not significantly different, have the highest AUC levels of all groups (Fig 1F). This is further confirmed by evaluating the mean RNAemia days, where it was observed that CD8-depleted animals had the longest RNAemia (Fig 1G). This suggests that early viremia and peak control associates with a late viremia resolution in CD4-competent but CD8-depleted animals, indicating that CD8+ T cells may not play a crucial role controlling early viremia but do so in mid to late viral clearance.

Taken together, these results confirm the requirement of CD4+ T cells for an optimal control of DENV2 replication during a primary infection. In addition, our findings propose an important role for CD4+ T cells controlling secondary ZIKV infection and that role being independent of prior DENV experience. Lastly, this indicates that CD8+ T cells may not be required for early ZIKV viral replication control in individuals with prior DENV2 immunity but are necessary for mid and late viral clearance.

### CD4^+^ T cell depletion results in higher detection of DENV NS1

To confirm viremia and to explore the role of CD4+ T cells in the generation of NS1 antigen, we measured DENV NS1 levels in serum. Since NS1 antigen is generally detected during days 1 to 9 p.i., we tested samples from baseline and days 3, 5, 7, 10, 15 and 30 post DENV2 infection (SFig 3). As expected, no detectable levels of NS1 were observed at day 0 on any of the groups. On days 3, 5, 7 and 10 p.i., an increase in NS1 levels was observed on all animal groups, with CD4-depleted animals having slightly higher NS1 levels in comparison to the undepleted animals by day 10 p.i.. By day 15 p.i., NS1 levels were significantly higher in depleted animals (p< 0.0001) compared to undepleted animals, where NS1 levels are completely undetectable. This is of particular interest because by this time, detection of DENV-NS1 is not expected. This suggests that the lack of CD4^+^ T cells enhances NS1 circulation, which correlates with enhanced pathogenesis, further confirming a protective role for CD4^+^ T cells during primary DENV infection.

### Lack of CD4+ T cells during a primary DENV2 infection does not hinder antibody binding to ZIKV in a secondary exposure

We evaluated the impact of CD4^+^ and CD8^+^ T cell depletion on the quantity and quality of the humoral response against DENV2 and ZIKV infections. Anti-DENV IgM and IgG levels were measured after primary DENV2 infection (Fig 2). CD4-depleted animals had significantly lower levels of anti-DENV IgM in comparison to the undepleted animals by days 10 and 15 p.i. (p< 0.001) (Fig 2A). Furthermore, total anti-DENV IgM produced from day 0 to day 15 p.i. was significantly lower in the CD4-depleted group in comparison to the control group (p= 0.007) (Fig 2C). On the other hand, anti-DENV IgG levels were statistically different between groups only on day 15 p.i., where the CD4-depleted group had lower levels compared to the control group (p< 0.001) (Fig 2B). Similar results were observed with the total anti-DENV IgG produced (from day 0 to day 30), where CD4-depleted animals had significantly lower IgG values in comparison to undepleted animals (p= 0.0153) (Fig 2D). IgG levels by days 60 and 90 p.i. were at similar levels in both groups. These results suggest a delay in B cell activation and in isotype class switching during primary DENV infection related to the lack of CD4^+^ T cells.

**Fig 2.**
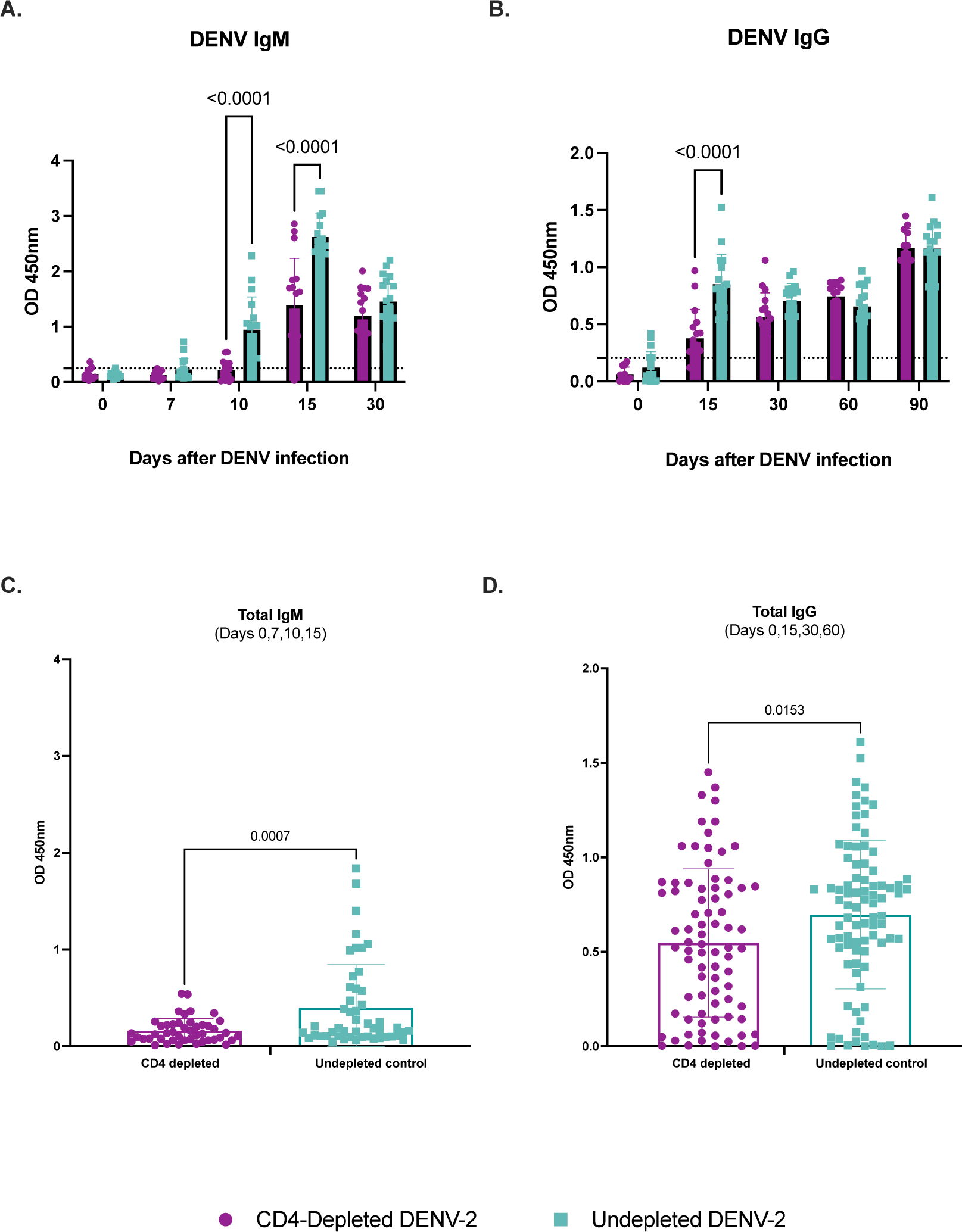
T cell depletion modifies serological profile during primary DENV2 infection. IgM and IgG response after DENV-2 infection was assessed using commercial ELISA tests. CD4-depleted animals are depicted in purple and undepleted animals are depicted in turquoise. Dotted lines indicate the limit of detection for each test. **(A-B)** Binding capacity of DENV IgM and IgG antibodies from CD4-depleted and undepleted animals after DENV-2 infection. **(C-D)** Total DENV IgM and IgG levels throughout different timepoints. Statistically significant differences among and within groups were calculated by two-way ANOVA using Tukey’s multiple comparisons test and unpaired t-tests.

Additionally, DENV and ZIKV binding IgM and IgG levels were measured after secondary ZIKV challenge (Fig 3). As expected, all animals had no detectable anti-DENV IgM at baseline and remain consistently low throughout days 7,10 and 15 p.c. (Fig 3A). At day 15 p.c. only one animal from the control group and one from the CD4+/DV2/CD8-/ZV group had values over the assay threshold, further indicating the positive role of CD4^+^ T cells in the magnitude of the humoral immune response during primary DENV infection. In contrast, all animals had similar detectable anti-DENV IgG levels at baseline (Fig 3B). As expected, ZIKV infection transiently boosted cross-reacting binding Abs which declined by day 30 p.c. in all groups (Fig 3B). However, the magnitude of this boost was significantly impacted on day 15 p.c. by the lack of CD4+ T cells. Lower anti-DENV IgG levels were observed in the two groups lacking CD4+ T cells before ZIKV challenge (double CD4-depleted and CD4+/DV2/CD4-/ZV). The significance between group CD4-/DV2/CD4-/ZV with groups CD4+/DV2/CD4+/ZV, CD4+/DV2/CD8-/ZV and CD4-/DV2/CD4+/ZV was statistically significant (p< 0.019, p< 0.008 and p< 0.047 respectively). The differences between the CD4+/DV2/CD4-/ZV with the CD4+/DV2/CD4+/ZV and CD4+/DV2/CD8-/ZV groups were also significant (p< 0.018 and p< 0.040 respectively). There were no differences among the three groups with CD4^+^ T cells before ZIKV infection. Altogether, these results indicate that DENV-experimented CD4^+^ T cells do not contribute to an optimal humoral immune response to ZIKV. This also suggests that DENV MBCs have a limited expansion during a secondary ZIKV infection and if they have, the expansion is CD4-independent. Anti-ZIKV IgM levels became detectable by day 7 p.c. with a peak by day 15 and started declining by day 30 p.c. (Fig 3C). By day 10 p.c., the control group had significantly higher levels than at least two groups, the doubled-depleted group (p< 0.030) and the CD4-/DV2/CD4+/ZV group (p< 0.033). By day 15 p.c., the differences between the control undepleted group and the two other groups undepleted of CD4+ T cells before ZIKV challenge were significant (CD4-/DV2/CD4+/ZV, p< 0.022 and CD4+/DV2/CD8-/ZV, p< 0.028). However, the two groups depleted of CD4+ T cells before ZIKV challenge have the lowest levels of ZIKV-binding IgM compared to the three undepleted groups (group CD4+/DV2/CD4+/ZV vs. CD4-/DV2/CD4-/ZV and CD4+/DV2/CD4-/ZV p< 0.0001 and p< 0.0001 respectively, group CD4-/DV2/CD4+/ZV vs. CD4-/DV2/CD4-/ZV and CD4+/DV2/CD4-/ZV p< 0.001 and p< 0.053 respectively and group CD4+/DV2/CD8-/ZV vs. CD4-/DV2/CD4-/ZV and CD4+/DV2/CD4-/ZV p< 0.001 and p< 0.004 respectively).

**Fig 3.**
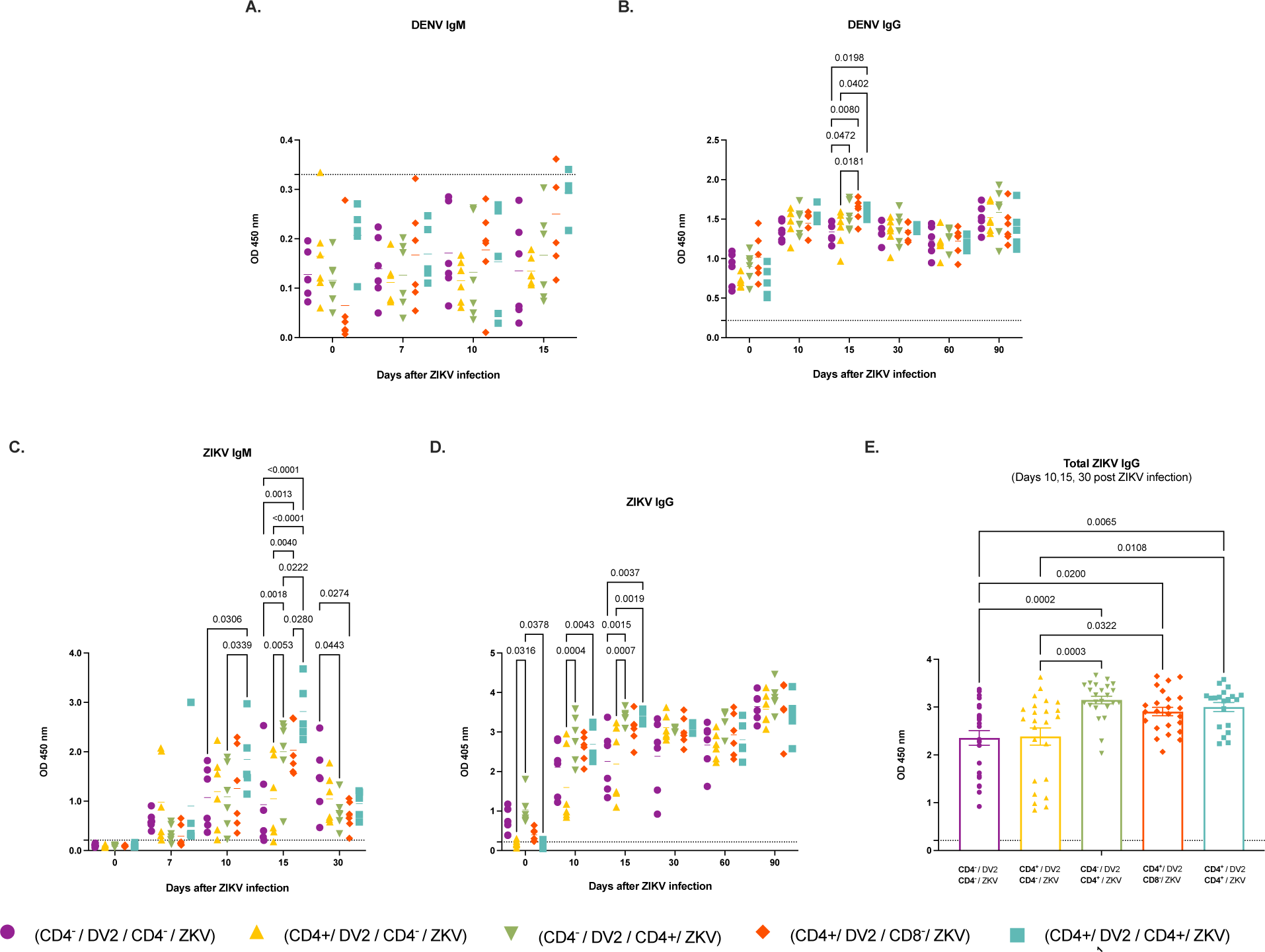
T cell depletion modifies serological profile during secondary heterologous ZIKV infection. IgM and IgG response after ZIKV infection in the presence of DENV immunity was assessed using commercial ELISA tests. Animal cohorts are depicted as follows based on treatment regime: CD4+/DV2/CD4+/ZV (shown in turquoise), CD4-/DV2/CD4+/ZV (shown in green), CD4+/DV2/CD4-/ZV (shown in yellow), CD4-/DV2/CD4-/ZV (shown in purple), and CD4+/DV2/CD8-/ZV (shown in red). Dotted lines indicate the limit of detection for each test. **(A-B)** Binding capacity of DENV IgM and IgG antibodies from CD4-depleted and undepleted animals after ZIKV infection. **(C-D)** Binding capacity of ZIKV IgM and IgG antibodies from CD4-depleted and undepleted animals after ZIKV infection. **(E)** Total ZIKV IgG levels after ZIKV infection were analyzed. Statistically significant differences among and within groups were calculated by two-way ANOVA using Tukey’s multiple comparisons test and unpaired t-tests, and by One-way ANOVA using Tukey’s multiple comparisons test.

Of note, some of the CD4-/DV2/CD4-/ZV animals had a late modest boost in ZIKV IgM levels by day 30 p.c. after the initial observed delay; interestingly, all depleted animals before ZIKV challenge induced the lowest IgM levels. Together, these observations suggest a delayed ZIKV-naïve B cell expansion in absence of CD4+ T cells.

Results from ZIKV-binding IgG Abs were similar to the IgM results (Fig 3D). By day 10 p.c. the two groups depleted of CD4+ T cells before the challenge showed the lowest values. The values were significantly lower in the group depleted of CD4+ T cells only before ZIKV challenge (CD4+/DV2/CD4-/ZV) compared to the CD4+/DV2/CD4+/ZV (p< 0.004) and CD4-/DV2/CD4+/ZV (p< 0.0001) groups. By day 15 p.c. this trend was maintained. Results reached statistical significance between the control undepleted group and the double-depleted group (p< 0.003) and the CD4+/DV2/CD4-/ZV group (p< 0.001) and between the CD4-/DV2/CD4+/ZV group with the CD4-/DV2/CD4-/ZV and the CD4+/DV2/CD4-/ZV groups (p< 0.001 and p< 0.0007 respectively).

An interesting finding was the presence of detectable cross-reacting anti-ZIKV IgG levels at baseline only in the group depleted of CD4+ T cells before primary DENV infection but not before ZIKV challenge (CD4-/DV2/CD4+/ZV) with significant higher values when compared to the CD4+/DV2/CD4-/ZV and the CD4+/DV2/CD4+/ZV groups (p< 0.031 and p< 0.037 respectively). The double-depleted group also had higher cross-reactive IgG levels compared to the undepleted groups but did not reach statistical significance. This result suggests an increased expansion of DENV-cross-reactive B cell clones in animals without CD4^+^ T cells during primary DENV infection.

To confirm our results on the anti-ZIKV IgG, we analyzed the total ZIKV-specific IgG produced on days 10, 15 and 30 p.c. (Fig 3E). Double-depleted animals had significantly less total ZIKV IgG production compared to the CD4-competent animals before ZIKV challenge (p= 0.0002 compared to CD4-/DV2/CD4+/ZV group, p= 0.0200 compared to CD4+/DV2/CD8-/ZV group and p= 0.0065 compared to CD4+/DV2/CD4+/ZV). Similarly, the CD4+/DV2/CD4-/ZV group had lower IgG production compared to the CD4-/DV2/CD4+/ZV, CD4+/DV2/CD8-/ZV and CD4+/DV2/CD4+/ZV groups (p= 0.0003, p= 0.0322 and p= 0.0108, respectively). In general, CD4-depleted animals before ZIKV challenge produced significantly less total ZIKV-specific IgG.

These results indicate that while CD4^+^ T cells during a primary DENV infection have limited impact in the binding of ZIKV-specific Abs, their absence may result in a significant induction of flavivirus cross-reactive B cell clones (Fig 2D) during a secondary infection. This also indicates that DENV-induced MBC populations generated during a primary DENV infection have limited contribution in the anti-ZIKV-binding Abs after ZIKV infection. In addition, these results suggest that CD4^+^ T cells are required for a more effective and specific antibody response against ZIKV.

### Recall memory to DENV NS1 is expanded by lack of CD4+ T cells before DENV2 infection

Next, we performed an anti-DENV NS1 assay using samples from baseline and days 15, 30 and 90 after primary DENV2 infection (SFig 4A). No DENV NS1 anti-IgG levels were detected on day 0 as expected. However, by day 15 p.i., depleted animals had significantly lower levels in comparison to the undepleted animals, similar to anti-DENV IgG levels previously presented (p= 0.0003). By day 30 p.i., both groups had reached similar anti-NS1 IgG levels. Interestingly, by day 90 p.i. depleted animals showed significantly higher anti-NS1 IgG levels compared to undepleted animals (p= 0.0003), altogether supporting the delay in Abs switch in this group as seen in the total IgG Abs response.

We also measured DENV NS1 anti-IgG levels in samples from baseline and days 15, 30 and 90 after secondary ZIKV challenge (SFig 4B). Of note, on baseline the group depleted of CD4^+^ T cells before DENV infection but not ZIKV (CD4-/DV2/CD4+/ZV) had higher values when compared to the groups with CD4^+^ T cells prior to DENV infection, and they were significantly higher particularly in comparison to the CD4+/DV2/CD4-/ZV group (p= 0.002). Although not significant, the double-depleted group had similarly high values in comparison to the groups with CD4+ T cells prior to DENV infection. This implies that the lack of CD4^+^ T cells before primary DENV infection also results in an increased durability of NS1 Abs. By day 15 p.c., we observed significantly higher values of DENV NS1 anti-IgG in the CD4-/DV2/CD4+/ZV group in comparison to all other groups (CD4-/DV2/CD4-/ZV, p= 0.0124; CD4+/DV2/CD4-/ZV, p= 0.0002; CD4+/DV2/CD4+/ZV, p= 0.0094 and CD4+/DV2/CD8-/ZV, p= 0.0152). This trend is maintained by day 30 p.c. (CD4-/DV2/CD4-/ZV, p= 0.0053; CD4+/DV2/CD4-/ZV, p< 0.0001; CD4+/DV2/CD4+/ZV, p< 0.0001 and CD4+/DV2/CD8-/ZV, p< 0.0001). Intriguingly, similar to what we stated on the previous section, these results seem to imply that deficiency of CD4^+^ T cells during the primary DENV2 infection modify the recall memory quantitatively resulting in an increased DENV NS1-cross-reactivivity induced by ZIKV.

Lastly, no significant differences between groups were observed on day 90 p.c.. Of note, some animals belonging to the CD4+/DV2/CD4-/ZV group exhibited a marked separation between IgG values, indicating that antibody repertoire may be different between individuals even sharing similar conditions.

### Depletion of CD4+ T cells only prior to DENV infection increases ZIKV NS1-binding IgG after ZIKV infection

To further study the impact of CD4+ T cells in the generation of cross-reactive and specific Abs, we measured anti-ZIKV NS1 IgG levels in serum at days 0, 15, 30 and 90 p.c. (SFig 5). Contrary to the total IgG, no animals had, regardless of their CD4+ T cell status, detectable anti-ZIKV-NS1 IgG Abs at baseline, indicating that those Abs are very specific. However, by day 15 p.c., only animals with CD4+ T cells before ZIKV challenge produced detectable IgG levels, with the exception of one animal from the CD4+/DV2/CD4-/ZV group (SFig 5). Double-depleted animals had the lowest IgG levels of all groups, significantly different from the control group (p= 0.0003) and the CD4-/DV2/CD4+/ZV group (p= 0.0004) on day 15 p.c. Additionally, NS1 IgG levels from the double-depleted group were significantly lower in comparison to the CD4-/DV2/CD4+/ZV group (p= 0.0071) also on day 30 p.c.. The latter group was consistently the highest IgG-producing group, even higher than the control group. Unsurprisingly, CD8-depleted animals behaved similarly to the control group as they both have intact CD4 T cells populations before the primary infection and the challenge. By day 90 p.c. almost all animals, regardless of their depletion status, had no NS1-IgG levels. Collectively, these data suggests that DENV-experimented CD4^+^ T cells have limited or no effect on the production of anti-ZIKV NS1-abs during a secondary ZIKV exposure.

### Early DENV2 neutralization activity is delayed by lack of CD4+ T cells

To determine the contribution of Abs to DENV2 and ZIKV neutralization and the impact of CD4+ and CD8+ T cell depletion, all groups were tested using FRNT and PRNT assays against DENV2 and ZIKV respectively (Fig. 2 and 3). Neutralization assays were completed for baseline and days 15, 30 and 90 p.i. for DENV-2 and days 15, 30 and 90 p.c. for ZIKV. The 50% effective concentration (EC) of neutralizing Abs is shown in Figures 3 and 4 as well. As expected, no animals had nAbs against DENV2 at baseline before primary infection (Fig 4A). By day 15 p.i., a significantly lower neutralizing activity is observed in the CD4-depleted animals in comparison to the undepleted group (p< 0.0001) (Fig 4B). This trend is not observed on day 30 p.i., when both groups behave similarly (Fig 4C). Endpoint titers dilutions are shown in Figures 3D and 3E in order to facilitate a better interpretation. Similar to the scenario described above, significantly lower EC50 values are observed in the depleted animals (p= 0.0032) (Fig 4D).

**Fig 4.**
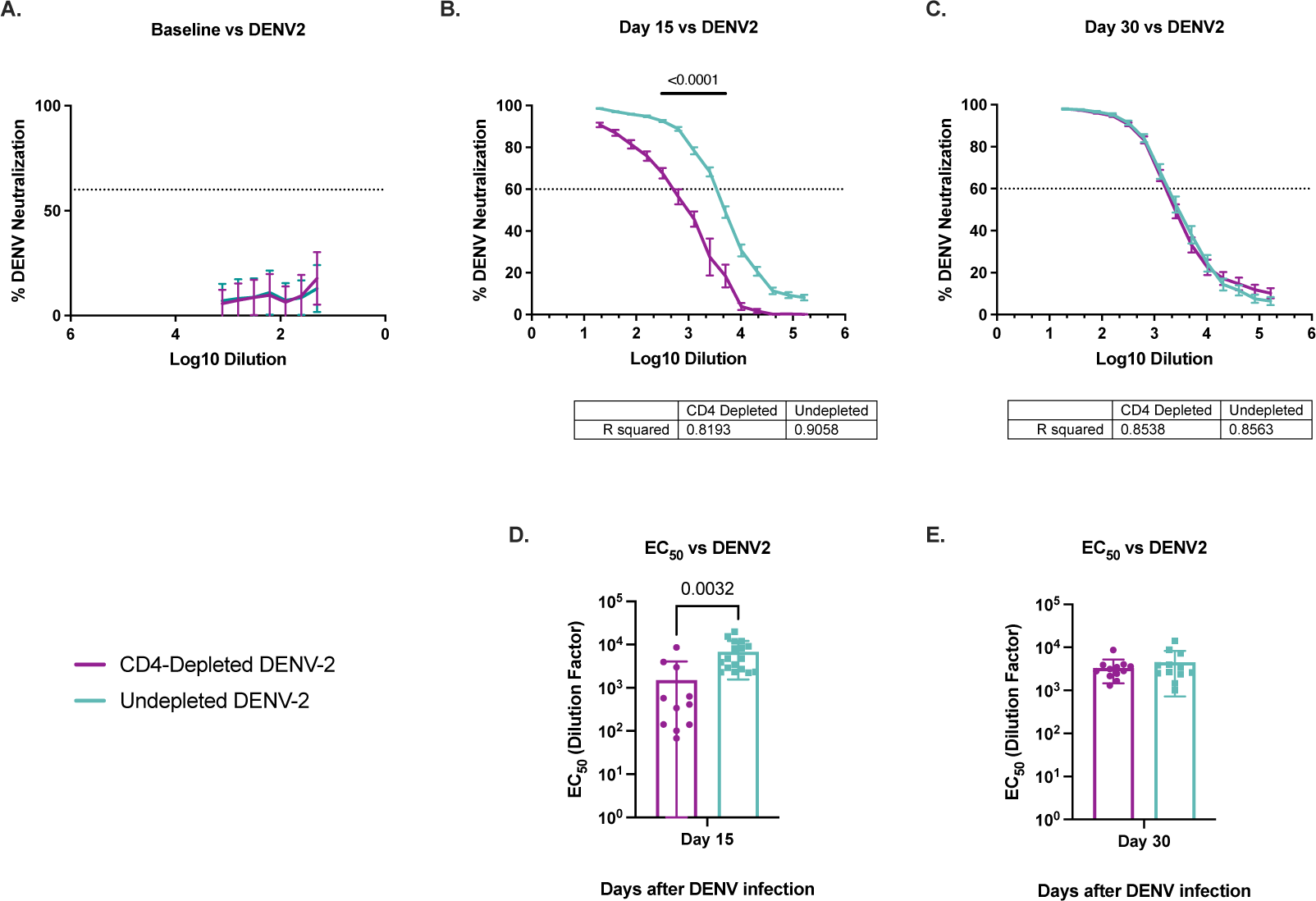
FRNT values of neutralizing antibodies against DENV-2 in depleted or undepleted flavivirus-naïve macaques. Neutralization capacity against DENV-2 is shown. CD4-depleted animals are depicted in purple and undepleted animals are depicted in turquoise. Dotted lines indicate the limit of detection for each test. **(A-C)** FRNT60 values of neutralizing antibodies against DENV-2 after DENV-2 infection are shown. **(D-E)** EC50 values of neutralizing antibodies against ZIKV after ZIKV infection. Statistically significant differences among groups were calculated by one-way and two-way ANOVA using the Tukey’s multiple comparisons test and unpaired t-tests.

PRNT assays were performed after ZIKV challenge; despite the detection of anti-ZIKV IgG at baseline in some groups, no neutralizing activity was observed at that time point against ZIKV (Fig 5A). However, on day 15 p.c., the CD4-double depleted group was the lowest at neutralizing ZIKV (Fig 5B). Animals belonging to the CD4-/DV2/CD4+/ZV group and interestingly those from group CD4+/DV2/CD8-/ZV, having a higher frequency of CD4+ T cells, have better neutralizing activity by day 15 p.c. compared to the other three groups – even the undepleted control group. Those differences were statistically significant compared to the double-depleted group (p= 0.0231 and p= 0.0155 respectively). Taken together, these results lead us to consider that DENV-primed CD4^+^ T cells have a limited contribution to an effective early neutralization during a secondary ZIKV infection and that the magnitude of neutralization can correlate with the frequency of those cells.

**Fig 5.**
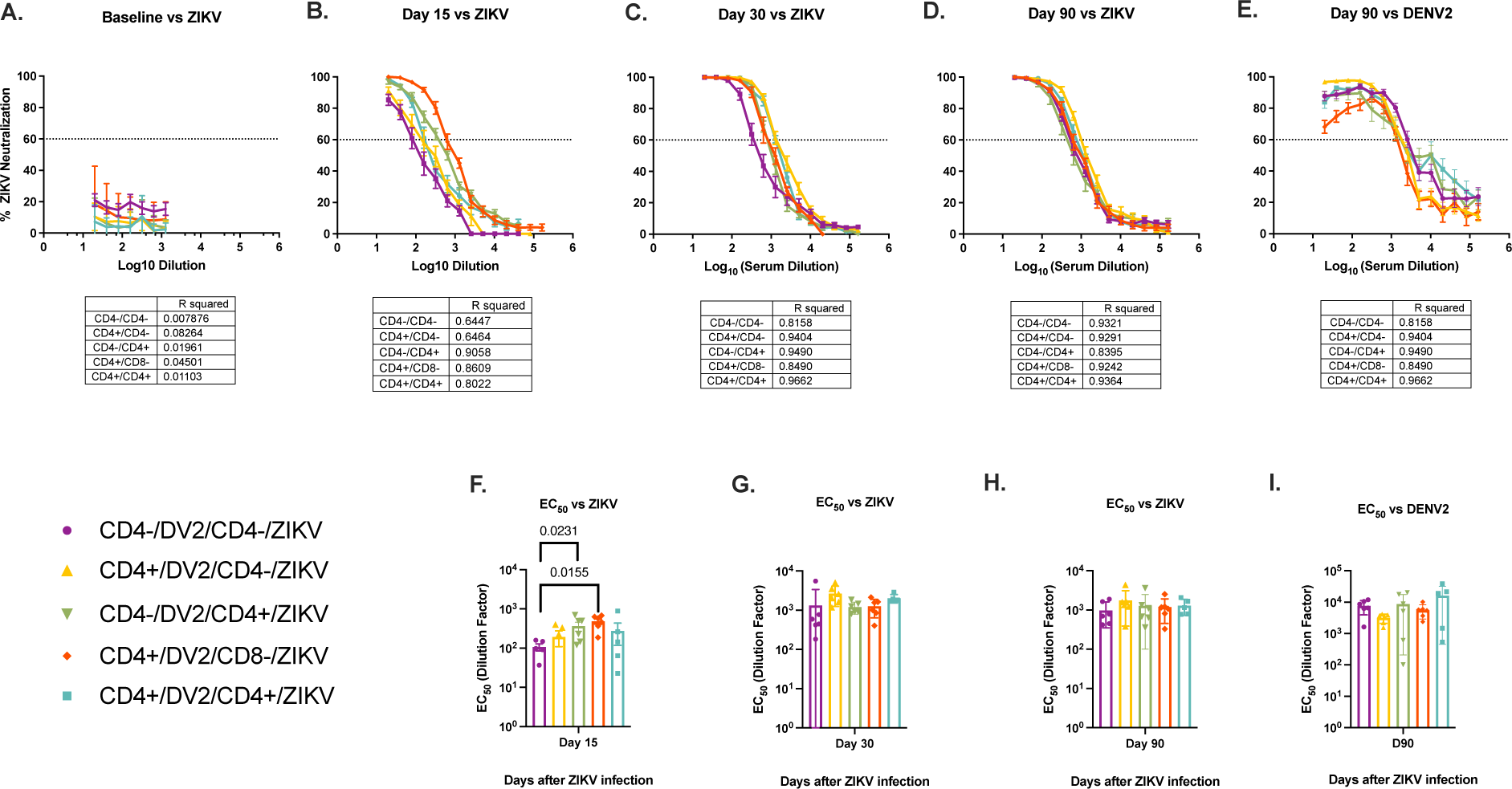
PRNT values of neutralizing antibodies against ZIKV in depleted or undepleted DENV-immune macaques. Neutralization capacity against ZIKV is shown. Animal cohorts are depicted as follows based on treatment regime: Animal cohorts are depicted as follows based on treatment regime: CD4+/DV2/CD4+/ZV (shown in turquoise), CD4-/DV2/CD4+/ZV (shown in green), CD4+/DV2/CD4-/ZV (shown in yellow), CD4-/DV2/CD4-/ZV (shown in purple), and CD4+/DV2/CD8-/ZV (shown in red). Dotted lines indicate the limit of detection for each test. **(A-D)** FRNT60 values of neutralizing antibodies against ZIKV after ZIKV infection are shown. **(E-G)** EC50 values of neutralizing antibodies against ZIKV after ZIKV infection. Significant differences among groups were calculated by one-way and two-way ANOVA using the Tukey’s multiple comparisons test and unpaired t-tests.

By day 30 p.c., a change in neutralization profile is observed in all groups except the double-depleted animals, which still show a delay in neutralization in comparison to the other groups (Fig 5C). By day 90 p.c., all groups behave similarly in terms of neutralization magnitude (Fig 5D). Endpoint titers dilutions are shown in Figures 4F-H in order to facilitate a better interpretation.

To further study the impact of CD4+ T cells on the humoral response, neutralization assays were performed against DENV2 using serum samples from 90 days post ZIKV challenge (Fig 5E and 5I). No statistical differences were detected between any of the groups.

### Lack of CD4+ T cells during a primary DENV2 infection compromises antibody specificity in a secondary ZIKV exposure

To further explore the role of CD4+ T cells in shaping the immune response against DENV and ZIKV, epitope reactivity against EDIII and NS1 was evaluated using a Luminex assay. EDIII was selected as a major neutralizing target and NS1 due to its proposed role in pathogenesis. Type-specific and cross-reactive responses were measured at day 90 p.i. after DENV2 and ZIKV infections against EDIII and NS1 antigens from all four DENV serotypes and ZIKV. To normalize the data a cross-reactivity index (CRI) was calculated by subtracting the background from the homologous value (values against ZIKV) and the heterologous values (values against DENV2) followed by dividing the subtracted homologous by the subtracted heterologous values (Fig 6).

**Fig 6.**
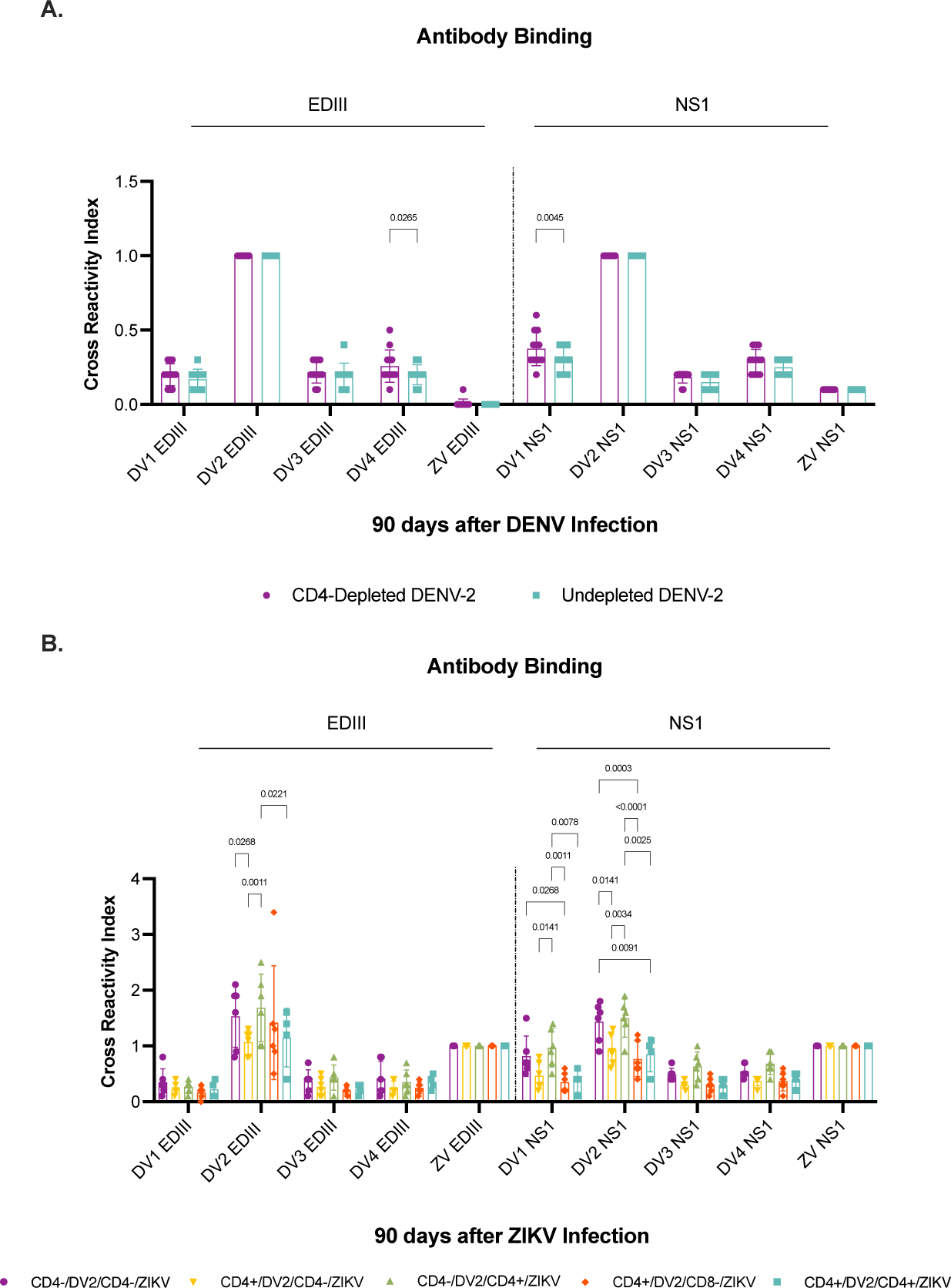
Epitope reactivity against DENV and ZIKV antigens in depleted and undepleted macaques. Antibody type-specific and cross-reactive responses against EDIII and NS1 antigens were assessed. Data generated by Luminex assay was normalized to a cross-reactivity index (CRI) by subtracting background (BSA) from the homologous value (values against infecting virus, HOM) and the heterologous values (values against heterologous virus, HET) followed by dividing the subtracted homologous by the subtracted heterologous values [<u>^(”#$%&’()^</u>]. **Upper panel.** CD4- (”*+%&’() depleted animals are depicted in purple and undepleted animals are depicted in turquoise. **(A)** Antibody response to EDIII (left side) and NS1 (right side) antigens from all four DENV serotypes and ZIKV 90 days after DENV2 infection**. Lower panel.** Animal cohorts are depicted as follows based on treatment regime: CD4+/DV2/CD4+/ZV (shown in turquoise), CD4-/DV2/CD4+/ZV (shown in green), CD4+/DV2/CD4-/ZV (shown in yellow), CD4-/DV2/CD4-/ZV (shown in purple), and CD4+/DV2/CD8-/ZV (shown in red). **(B)** Antibody response to EDIII (left side) and NS1 (right side) antigens from all four DENV serotypes and ZIKV 90 days after ZIKV infection. Significant differences among groups were calculated by two-way ANOVA using the Tukey’s multiple comparisons test.

As expected, after primary DENV2 infection, both undepleted and depleted animals had a high similar type-specific response against EDIII and NS1 to the infecting serotype DENV2, and a similar limited cross-reactivity against EDIII to the other DENV serotypes, with a significant difference observed in DENV4 EDIII (p= 0.0265) (Fig 6A). As reported in the literature, primary DENV infection in flavivirus-naïve individuals triggers minimal or no cross-reactivity to ZIKV (Collins et al., 2017; Duffy et al., 2009). Both groups show a similar high type-specific response against DENV2 NS1 protein. In contrast to the response to EDIII domain, a higher cross-reactive response against NS1 to the other DENV serotypes, particularly against DENV serotype 1, was observed (p= 0.0045). Interestingly, there was a limited but detectable cross-reactivity against ZIKV NS1 protein in both groups (Fig 6A). This suggests that during a primary DENV infection, CD4+ T cells have a different weight in the responses against EDIII domain and NS1 protein.

A more complex scenario is observed after secondary ZIKV infection (Fig 6B). All animals show an increase in the type-specific response against EDIII and NS1 towards the infecting ZIKV, but their reactivity against other DENV serotypes varies. Additionally, all groups show reactivity against DENV2 antigens. Of note, two animals from the undepleted control group NS1 (MA254 and MA264) had a higher type-specific response against DENV2 EDIII, and two had a lower response (MA311 and MA350) (Fig 6B), indicating that antibody repertoire can be different between individuals, a point to take into consideration in vaccine design. Lastly, as expected, the CD4+/DV2/CD8-/ZV group behaved very similar to the CD4-competent groups, demonstrating that CD8^+^ T cell depletion does not have an impact in epitope reactivity (Fig 6B). Taken together, these results indicate that while CD8^+^ T cell depletion increases CD4^+^ T cell frequency and results in better ZIKV viremia control, quantity has no effect on the specificity of epitope recognition. Overall, the cross-reactivity to NS1 protein was more significant among DENV serotypes than to EDIII domain suggesting that, like after a primary DENV infection, CD4+ T cells have a different weight in the response against EDIII domain and NS1 protein after a secondary ZIKV infection in DENV-immune individuals. Animals belonging to all groups, regardless of depletion condition, show an increased but similar response against EDIII and NS1 to the infecting ZIKV. These results suggest that pre-existing DENV immunity has a very limited impact in the cross-reactivity to ZIKV EDIII domain and NS1 protein, and that CD4^+^ T cells have a limited role regulating that cross-reactivity to those ZIKV targets.

### CD4+ T cells modulate early response IgG+ MBC activation and functionality during a heterologous ZIKV infection with previous DENV-2 immunity

To determine the contribution of CD4+ T cells in modulating the surface IgG+ memory B cell (IgG MBC) (CD3-CD20+CD27+sIgG+) response to ZIKV, we measured expression levels of surface IgG in total MBCs via Flow Cytometry. Whole blood for Immunostaining was collected on baseline and days 7, 15, and 30 p.c. with ZIKV. By day 15 p.c., a discrete peak in frequencies was observed in all groups, except the CD4+/DV2/CD8-/ZV group which remain at the baseline levels (Fig 7A). On day 15 p.c., the CD4+/DV2/CD4+/ZV and the CD4-/DV2/CD4+/ZV were the only groups showing a significant increase in IgG expression when compared to their baseline status. By day 30 p.c., all groups showed similar levels of IgG expression to their baseline status. We did not detect a significant difference in IgG expression among groups compared at each timepoint suggesting that sIgG expression is not impacted by depletion treatment status. Taken together, this data suggests that a secondary ZIKV infection in individuals with prior DENV-2 immunity, IgG+ MBCs levels suffer limited increase in frequency and if it happens it is CD4+ T cell-dependent.

**Fig 7.**
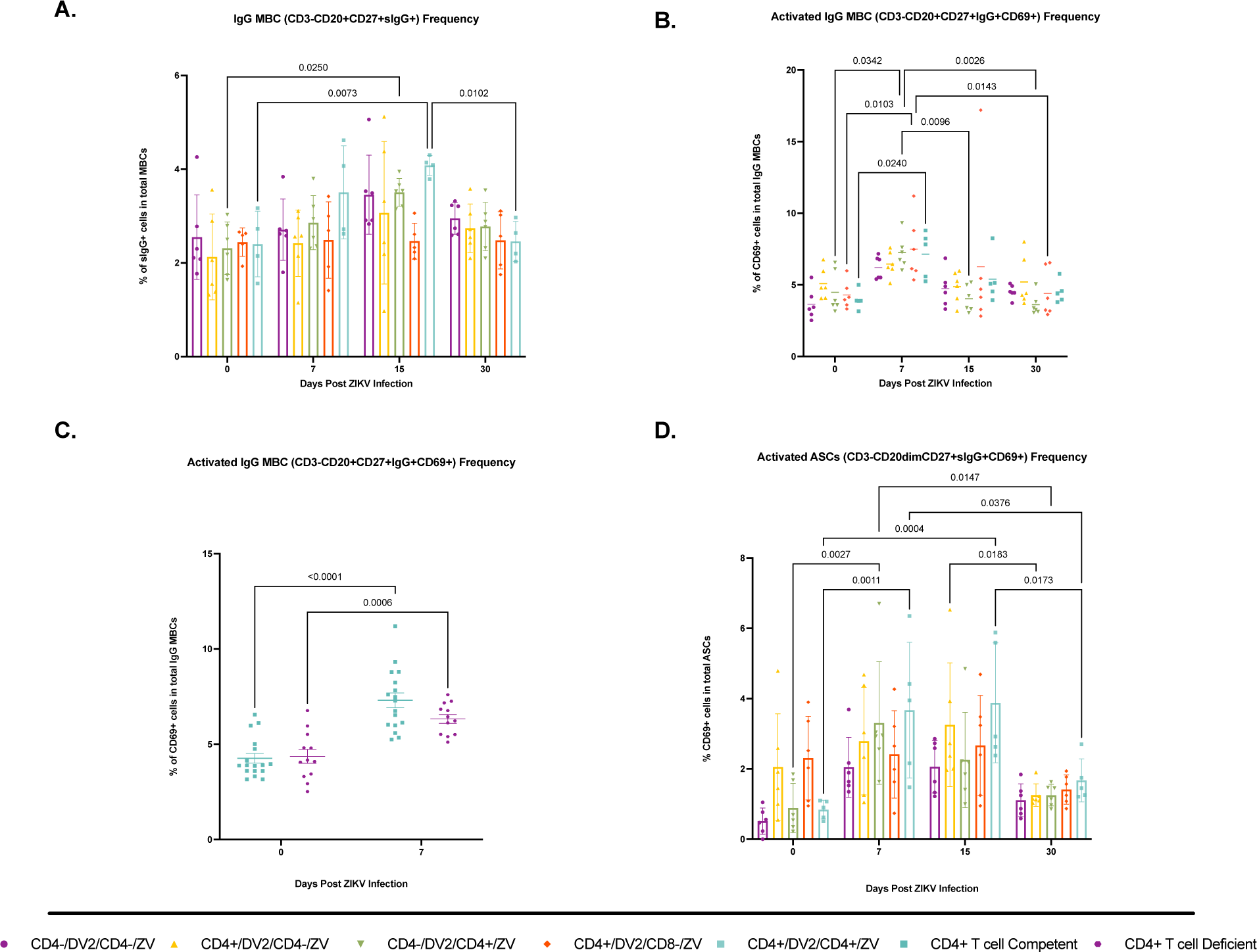
IgG memory B cell and ASC population kinetics after ZIKV infection in CD4+ T cell-competent and -deficient animals with previous DENV-2 immunity. The frequency of IgG MBCs and ASCs was assessed by immunophenotyping using flow cytometry. Animal cohorts are depicted as follows based on treatment regime: CD4+/DV2/CD4+/ZV (shown in turquoise), CD4-/DV2/CD4+/ZV (shown in green), CD4+/DV2/CD4-/ZV (shown in yellow), CD4-/DV2/CD4-/ZV (shown in purple), and CD4+/DV2/CD8-/ZV (shown in red). **(A)** Percentages of surface IgG expression in total MBCs (CD3-CD20+CD27+sIgG+). **(B)** Frequencies of CD69+ cells in total IgG+ MBCs (CD3-CD20+CD27+sIgG+CD69+). **(C)** Frequencies of CD69+ cells in total IgG MBCs with all cohorts organized into CD4+ T cell competent (shown in turquoise) and CD4+ T cell depleted (shown in purple). **(D)** Frequencies of CD69+ cells in total ASCs (CD3-CD20dimCD27+CD69+). Statistically significant differences among groups were calculated by two-way ANOVA using the Tukey’s multiple comparisons test and unpaired t-tests.

To determine if CD4+ T cells have a role in modulating the activation pathways of IgG MBCs after the ZIKV challenge, we surveyed the expression of CD69 in this cell population. Compared to their baseline, a significant increase in the frequency of IgG MBCs expressing CD69 was observed at day 7 p.c. only on those groups with intact CD4+ T cells before ZIKV challenge (CD4-/DV2/CD4+/ZV, p< 0.034; CD4+/DV2/CD4+/ZV, p< 0.024; and CD4+/DV2/CD8-/ZV, p< 0.010) (Fig 7B and 7C). Taken together, this data suggests that IgG MBCs go through CD4^+^ T cell-dependent interactions for their activation during a secondary ZIKV infection.

To determine if CD4+ T cells have a role in the differentiation of activated B cells into Antibody Secreting Cells (ASCs) (possibly through Germinal Centers formation) during a heterologous ZIKV infection we measured CD69 expression in total ASCs (CD20^dim^CD3^-^CD27^+^). Despite our gating strategy being designed to characterize rhesus macaque plasmablast populations (Silveira et al., 2015) (Zhang et al., 2019), we were unable to segregate that population. However as previously reported by us (Marzan-Rivera et al., 2022), we idetified the total ASC population and observed a trend in activation of that population during the ZIKV infection. At day 7 p.c., we detected a significant increase in CD69 expression in the CD4+/DV2/CD4+/ZV and the CD4-/DV2/CD4+/ZV groups, compared to the other groups (Fig 7D, p< 0.005). This peak in CD69 expression was maintained up to day 15 p.c. in the CD4+/DV2/CD4+/ZV group followed by a significant drop in CD69 expression levels at day 30 p.c.. Levels of expression observed during this timepoint were of similar magnitude as on baseline. On the other hand, groups that lacked CD4+ T cells before the ZIKV infection (CD4+/DV2/CD4-/ZV and CD4-/DV2/CD4-/ZV) showed a trend of lower levels of ASCs expressing CD69 compared to the CD4^+^ T cell competent groups before the ZIKV infection. Our data points to a *de novo* B cell response pathway that is T cell-dependent to promote the formation of both IgG MBCs and ASC subsets with good neutralizing capacity towards ZIKV.

### CD4^+^ Peripheral T helper cells (pTfh) are present in circulating blood of Rhesus macaques during heterologous ZIKV infection with prior DENV2 immunity

To confirm the presence and frequency of pTfh CD4+ T cells, we measured expression levels of surface markers such as CXCR5, CXCR3, PD-1, and ICOS via flow cytometry analysis. Chemokine receptor CXCR3+ expression has been associated with Type 1 T helper immune responses (Th1) (Sánchez-Vargas and Mathew, 2019; Sandberg et al., 2021; White et al., 2021). Moreover, PD-1 and ICOS expression is associated with B cell interaction in order to induce B cell activation (Lei et al., 2021; Shi et al., 2018). This analysis was performed only on groups CD4+/DENV2/CD4+/ZIKV, CD4-/DENV2/CD4+/ZIKV and CD4+/DENV2/CD8-/ZIKV because they have CD4+ T cell populations before ZIKV infection. Frozen PBMCs were used for Immunostaining on day 0 (baseline), day 7, and day 15 p.c. with ZIKV. Surface expression of CD4+/CXCR5+ was detected at similar levels in all three tested groups, confirming circulation of pTfh cells. No significant differences were established among groups after ZIKV challenge on day 7 p.c.. By day 15 p.c. the control undepleted group and the group lacking CD4+ T cells before DENV showed a trend to higher frequencies compared to the group depleted of CD8+ T cells before ZIKV challenge, and that difference reached significant values between that group and the CD4-/DENV2/CD4+/ZIKV group on day 15 p.c. (p< 0.039) (Fig 8A). Despite having similar frequency of pTfh T cells (CD4+/CXCR5+), results demonstrate significance on days 0, 7, and 15 p.c. regarding the frequency of total pTfh TH1 CD4+ T (CXCR5+ and CXCR3+) among groups with the group depleted of CD4+ T cells before DENV infection, having significant lower frequencies of those cells since baseline (p< 0.000 and p= 0.0034) and on day 7 p.c. (p< 0.000 and p= 0.0306) compared to groups CD4+/DENV2/CD4+/ZIKV and CD4+/DENV2/CD8-/ZIKV, respectively. On day 15 p.c. there was a marked decrease in the frequency of TH1 cells in the CD4+/DENV2/CD8-/ZIKV group compared to the control undepleted group (p= 0.0009). The difference in frequency was also significant between the control group and the CD4-/DENV2/CD4+/ZIKV group (p= 0.0012) (Fig 8B). Further, we wanted to assess the co-expression of B cell co-stimulating markers PD-1 and ICOS in order to correlate these results with B cell activation. The frequency of total CD4+ pTfh T cells CD4+/CXCR5+/CXCR3+/PD-1+/ICOS+ showed a trend to increase from baseline to day 7 p.c. in all three groups. We found same profile as with the TH1 pTfh cells were the CD4-/DENV2/CD4+/ZIKV show statistically significantly lower frequency of these co-stimulating markers at baseline and on days 7 and 15 p.c. compared to groups CD4+/DENV2/CD4+/ZIKV and CD4+/DENV2/CD8-/ZIKV (p<0.0001 in all comparisons) (Fig 8C). Gating strategy is shown in Figure 6D.

**Fig 8.**
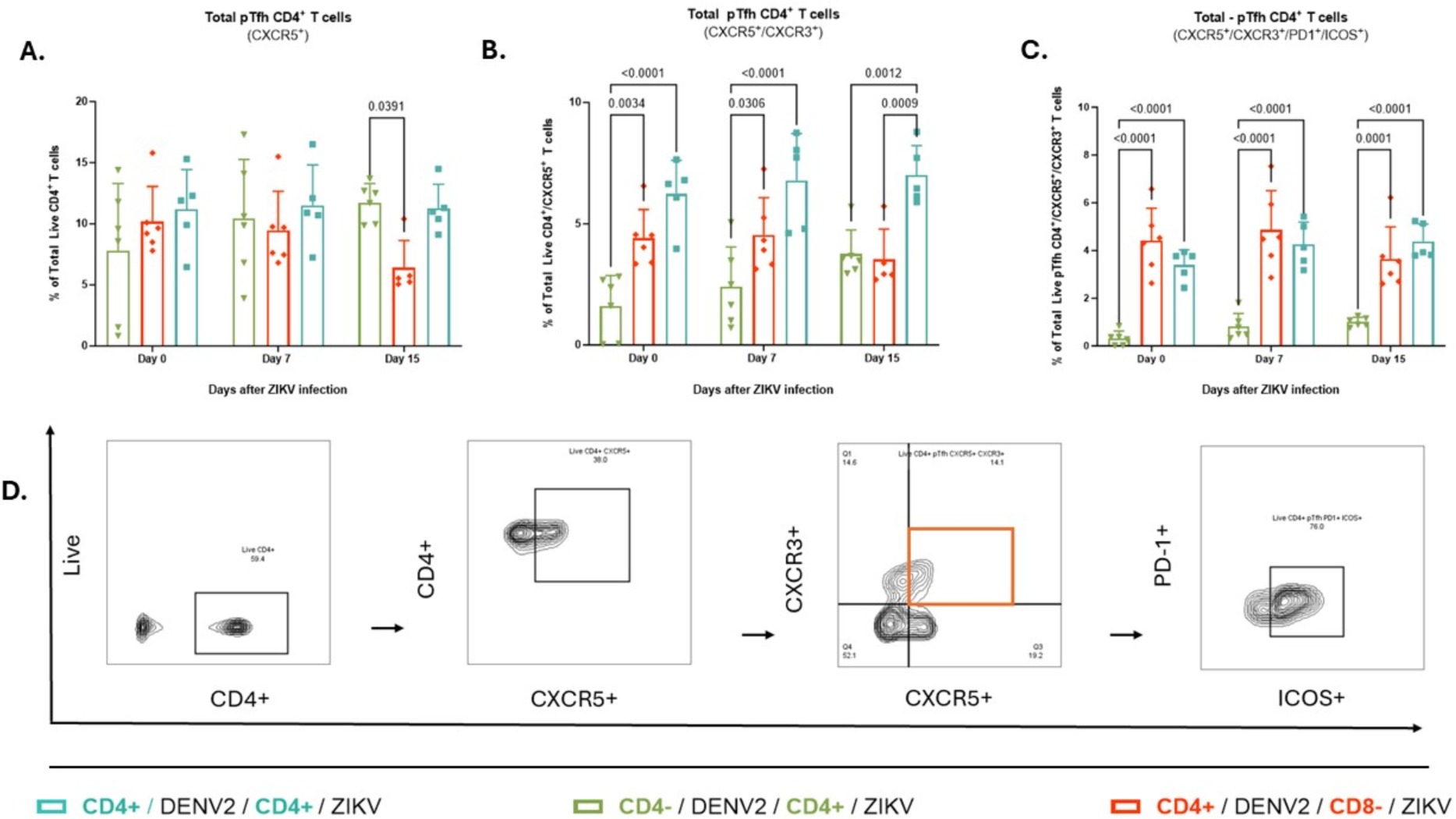
Total pTfh CD4+ T cell characterization. Immunophenotyping of CD4+ pTfh T cells was assessed via flow cytometry using frozen PBMCs collected on baseline, day 7, and day 15 pi with ZIKV. Animal cohorts are depicted as follows based on treatment regime: CD4+/DV2/CD4+/ZV (shown in turquoise), CD4-/DV2/CD4+/ZV (shown in green), and CD4+/DV2/CD8-/ZV (shown in red). **(A)** Frequency of total CD4+ T cells expressing follicular canonical marker CXCR5. **(B)** Frequency of total CD4-pTfh T cells expressing CXCR5 and CXCR3. **(C)** Frequency of total pTfh CD4 T cells expressing B cell co-stimulating surface makers PD1, and ICOS. **(D)** Gating Strategy for total pTfh CD4+ T cells characterization based on CXCR5, CXCR3, PD1, and ICOS surface markers expression. Comparisons between groups are reported as multiplicity-adjusted p values performed by two-way ANOVA using Tukey’s multiple comparisons tests and unpaired t-test.

When evaluating the total CD4+ T cell memory phenotype subsets (naïve CD28+CD95-, effector memory (EM) CD28-CD95-, and central memory (CM) CD28+CD95+), CM pTfh CD4+ T cells (CD4+/CD28+/CD95+) remain to be the more abundant phenotype of CD4+ memory T cells. (data not shown). To explore the impact of CD4+ and CD8+ T cell depletion on the frequencies of CM CD4+ pTfh T cells (CD4+/CD28+/CD95+/CXCR5+) we measured frozen Peripheral Blood Mononuclear Cells (PBMC) collected on day 0 (baseline), day 7, and day 15 p.c. with ZIKV via flow cytometry analysis (Fig 9A). Overall, there was a non-significant trend to increased frequency in all groups from day 0 to day 7 p.c. followed by a contraction on day 15 p.c. On baseline, the frequency was significantly lower in the CD4-/DENV2/CD4+/ZIKV group compared to the CD4+/DENV2/CD8-/ZIKV group (p< 0.002). However, there were no differences among groups after the challenge. Results also demonstrate a limited expansion of CM CD4+ pTfh T cell subsets co-expressing CD28+/CD95+/CXCR5+/CXCR3+ in all groups on day 7 p.c. with a contraction in the CD4+/DENV2/CD8-/ZIKV group by day 15 p.c.. As similarly seen with prior CD4+ pTfh profiles, the CD4+ T cell-deficient group before primary DENV infection result in the lowest frequency since baseline (p< 0.0001 and p= 0.0024), on day 7 (p< 0.0001 and p= 0.0209), and on day 15 p.c. (p< 0.0001 and p= 0.0008) compared to groups CD4+/DENV2/CD4+/ZIKV and CD4+/DENV2/CD8-/ZIKV, respectively (Fig 9B). We also measured the CM subpopulation co-expressing the B cell co-stimulating markers PD-1+ and ICOS+ (CD28+/CD95+/CXCR5+/CXCR3+/PD1+/ICOS+) (Fig 8C). All three groups showed the same trend as in the other CM population with a very limited expansion on day 7 and a trend to contraction on day 15 p.c. in the control undepleted and in the CD4+/DENV2/CD8-/ZIKV groups but not in the CD4-/DENV2/CD4+CD4+/ZIKV, which remains with similar frequency as day 7 p.c.. Interestingly, at baseline the CD4+/DENV2/CD8-/ZIKV group showed a frequency of these cells that is significantly higher compared to the control group (p< 0.035). But again, the CD4+ T cell-deficient group before primary DENV infection resulted in the lowest frequency since baseline (p= 0.0412 and p< 0.000), on day 7 (p= 0.0046 and p< 0.0001), and on day 15 p.c compared to groups CD4+/DENV2/CD4+/ZIKV and CD4+/DENV2/CD8-/ZIKV respectively (Fig 9C). There were no significant differences among groups on day 15 p.c. This data on the pTfh cells subsets show a detectable, but minimal expansion of the pTfh cells, and its CM compartment, during heterologous ZIKV infection in prior DENV2 immune individuals.

**Fig 9.**
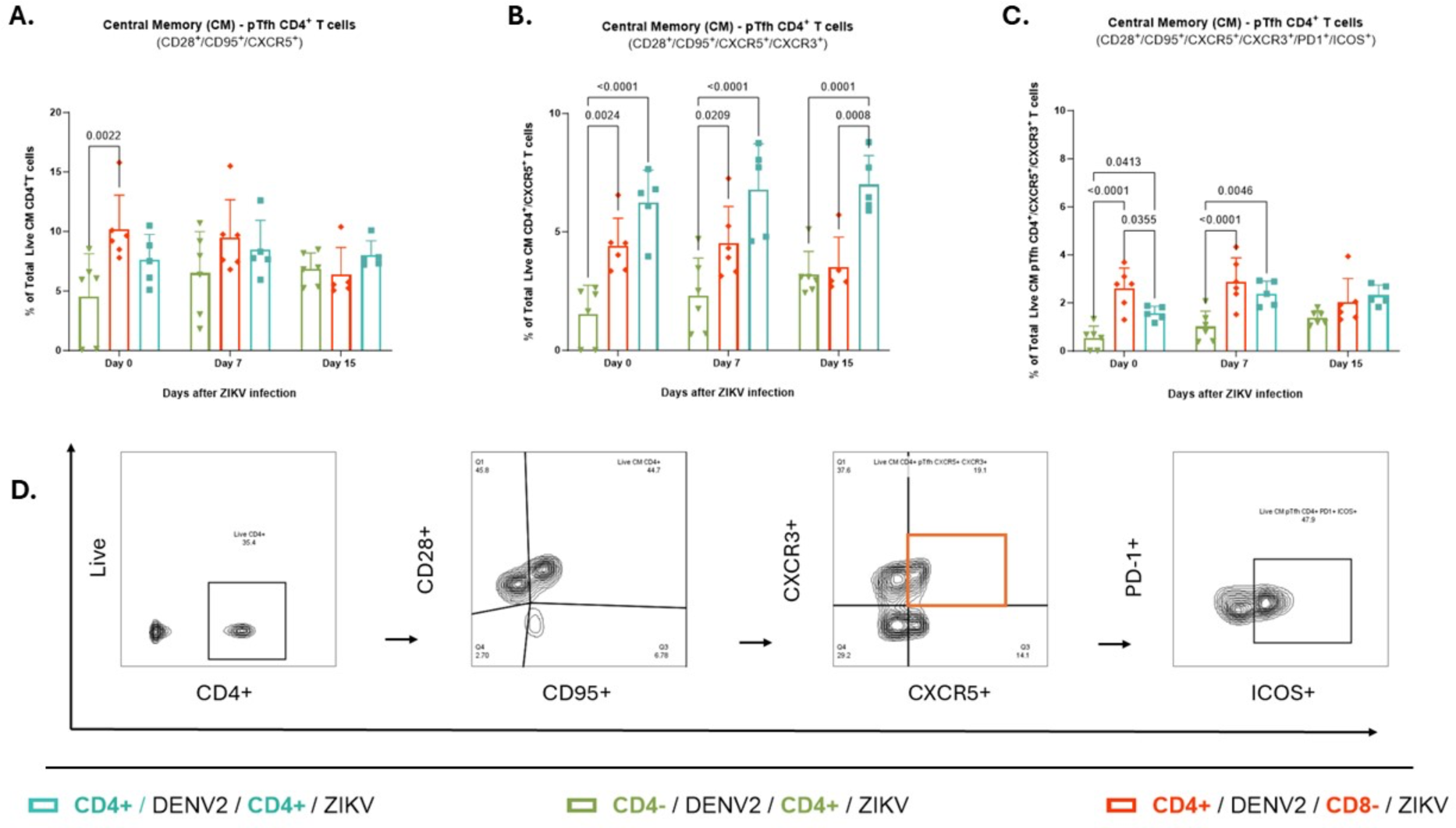
Total CM-pTfh CD4+ T cell characterization. Immunophenotyping of CM-pTfh CD4+ T cells after secondary ZIKV infection was accessed via flow cytometry using frozen PBMCs collected on baseline, day 7, and day 15 pi with ZIKV. Animal cohorts are depicted as follows based on treatment regime: CD4+/DV2/CD4+/ZV (shown in turquoise), CD4-/DV2/CD4+/ZV (shown in green), and CD4+/DV2/CD8-/ZV (shown in red). **(A)** Frequency of total CM CD4+ T cells expressing CXCR5. **(B)** Frequency of total CM-pTfh CD4+ T cells expressing CXCR5 and CXCR3. **(C)** Frequency of total CM-pTfh CD4+ T cells expressing B cell co-stimulating surface makers PD1, and ICOS. **(D)** Gating Strategy for total CM-pTfh CD4+ T cells characterization based on CXCR5, CXCR3, PD1, and ICOS surface markers expression. Comparisons between groups are reported as multiplicity-adjusted p values performed by two-way ANOVA using Tukey’s multiple comparisons tests and unpaired t-test.

### Total and DENV primed CM pTfh cells modulate both B cell subset activation and serum antibody response functionality towards ZIKV

To determine if the frequencies of peripheral pTfh cells are directly associated with the humoral response to the secondary infection with ZIKV, we performed Pearson correlation and simple linear regression analysis comparing pTfh cell frequencies with the neutralization assays and the IgG^+^ MBC immunophenotyping data. For the IgG^+^ MBC analysis, we compared three pTfh cell phenotypes: CXCR5^+^, CXCR5^+^CXCR3^+^, and CXCR5^+^CXCR3^+^PD1+ICOS^+^, at days 7 and 15 p.c., with frequencies of activated IgG^+^ MBCs, and activated ASCs, at 7 days p.c.. While we did perform correlation analysis comparing the pTfh cell frequencies with frequencies of total IgG^+^ MBCs, at 15 days p.c., we did not observe any significant correlation (data not shown). Interestingly, we observed a very low yet significant positive correlation (R^2^= 0.3237, P= 0.0214) when we compared frequencies of CXCR5^+^ pTfh cells from day 7 p.c.. with total IgG^+^ MBC frequencies from day 15 p.c. from all animals in the study (SFig 6A). From these observations, we decided to perform these correlation analyses with the experimental groups separately as we observed that correlations vary significantly depending on both cell phenotype and T-cell depletion treatment. When comparing the CXCR5^+^CXCR3^+^ pTfh cell phenotype, from Day 7 p.c., with the activated IgG^+^ MBC phenotype, we observed very high and significant correlations in both the CD4+/DV2/CD4+/ZV and the CD4-/DV2/CD4+/ZV groups (R^2^= 0.8790, P= 0.0186 and R^2^= 0.7424, P= 0.0274, respectively) (SFig 6B and 6C). In addition, a very positive and significant correlation with activated IgG^+^ MBC and PD1^+^ICOS^+^ pTfh cell frequencies was observed in the CD4-/DV2/CD4+/ZV group (R^2^= 0.7028, P= 0.0371) (SFig 6D). When comparing the CXCR5^+^ pTfh cells, from Day 7 p.c., with activated ASCs, we only observed a highly negative and significant correlation in the CD4-/DV2/CD4+/ZV group (R^2^= -0.7, P= 0.0378) (SFig 6E). On the other hand, we did not observe a significant correlation when performing this comparison in the CD4+/DV2/CD4+/ZV group. Furthermore, when we compared the frequencies of PD1^+^ICOS^+^ pTfh cell, from Day 7 p.c., with activated ASCs we observed a very high and significant positive correlation in the CD4+/DV2/CD4+/ZV group only (R^2^= 0.8657, P= 0.0218) (SFig 6F) These observations suggest that not only the B cell response is modulated by possible interactions with these pTfh cell subsets, but also these mechanisms are strongly influenced by the nature of the initial priming with DENV. Also, the opposed correlation (MBC vs. ASC) in the CD4-/DV2/CD4+/ZV group strongly suggest that lack of DENV-experimented pTfh modify the interactions with the MBC and ASC in different ways. As part of this analysis, we also measured possible correlations between these B cell subsets and these same pTfh cell subsets but focusing on CM pTfh cells. Here, we only observed very high and significant positive correlations between the CXCR5^+^CXCR3^+^ and PD1^+^ICOS^+^ CM pTfh cells, from day 7 p.c., and activated IgG^+^ MBCs and activated ASCs in the CD4+/DV2/CD4+/ZV group (SFig 7). This data adds new insight to the impact of the initial DENV priming in the secondary ZIKV infection response, as CM pTfh cells only show a significant correlation in the expansion of activated B cell subsets in animals with DENV-primed CD4^+^ T cells. Lastly, the CD4+/DV2/CD8-/ZV group showed no significant correlations between any of the pTfh, and B cell subsets analyzed (data not shown). This could suggest a role or possible interactions of CD8+ T cells with either pTfh cells and/or B cell subsets during the ZIKV infection.

To characterize the impact of pTfh cells in the functionality of the antibody response towards the ZIKV infection, we performed Pearson correlation and linear regression analysis comparing the previous pTfh cell phenotypes, during day 7 and 15 p.c.., with the EC50 data (from ZIKV and DENV neutralizations) gathered from days 30 and 90 p.c.. Similar to what we observed in the correlation analysis with IgG^+^ MBCs and ASCs, only the CD4+/DV2/CD4+/ZV group showed highly positive and significant correlations between pTfh cell phenotypes and the ZIKV EC50 values (SFig 8). When comparing frequencies of CXCR5^+^ pTfh cells, from both days 7 p.c. and 15 p.c., with EC50 values from day 30 p.c., we observed a high and significant positive correlation at both timepoints (R^2^= 0.8317, p= 0.0309 and R^2^= 0.9115, p= 0.0115, respectively) (SFig 8A and 8B). In the EC50 from day 90 p.c., we only observed significant positive correlations with the frequencies of PD1^+^ICOS^+^ pTfh cells in both day 7 and 15 p.c. timepoints (R^2^= 0.7796, p= 0.0472 and R^2^= 0.9182, p= 0.0102, respectively) (SFig 8C and 8D). We did not observe any significant correlations with CXCR5^+^CXCR3^+^ pTfh cells (data not shown). As part of this analysis, correlations were performed with CM pTfh cell subsets as well. Similar to previous results, only the CD4+/DV2/CD4+/ZV group showed significant positive correlations between CM pTfh cell frequencies and ZIKV EC50 values from days 30 and 90 p.c.. Interestingly, for both the CXCR5^+^ from day 15 p.c. and the CXCR5^+^CXCR3^+^ pTfh cells from days 7 and 15 p.c., a strong positive correlation was observed when compared with EC50 values from day 30 p.c. when comparing all animals in the study (R^2^= 0.3282, p= 0.0204, R^2^= 0.2863, p= 0.0269, R^2^= 0.5100, p= 0.0019 respectively) (SFig 8E, 8F and 8G). Similar to previous observations, there were significant positive correlations for CXCR5^+^ CM pTfh cells from days 7 and 15 p.c., when compared to EC50 values from day 30 p.c. (R^2^= 0.9505, p= 0.0470, and R^2^= 0.9063, p= 0.0125 respectfully) (SFig 8H and 8I). Lastly, the only positive correlation observed in EC50 values from day 90 p.c. was observed when comparing frequencies of PD1^+^ICOS^+^ CM pTfh cells from day 15 p.c. (R^2^= 0.8214, p= 0.0339) (SFig 8J). Taken together, this data adds new insight to the importance of the nature of the initial priming with DENV and its impact on the ZIKV infection, as we also saw strong correlations between CM pTfh cell phenotypes and EC50 values in the CD4+/DV2/CD4+/ZV group only. This suggests that DENV-primed CM pTfh cells may play an important role in modulating an efficient antibody response to ZIKV. In addition, this data adds more weight to the role of ICOS^+^PD1^+^ pTfh cells in priming a robust Abs response with efficient neutralizing capacity towards flavivirus. Interestingly, we observed no significant correlations when we compared total and CM pTfh cells with EC50 values, from day 90 p.c., that neutralized DENV-2 in any experimental group (data not shown). These results suggest that, while possible DENV-primed CM pTfh cells help to modulate the potency of nAbs to ZIKV, these mechanisms between pTfh cells and the B cell response may be only part of the type-specific humoral response to ZIKV. These results further support a T-cell dependent humoral immune response to ZIKV in DENV-experimented individuals. Very interesting and highly relevant is the fact that mechanistically the CD8+ T cell depletion abrogates the significant correlations between pTfh cell phenotypes and the ZIKV EC50 values in the CD4+/DV2/CD4+/ZV group. This strongly indicates a role of that cell population shaping the nAbs properties.

### Principal component analysis recapitulates infection dynamics

To reduce the dimensionality of our data – which included ELISAs, neutralization assays, flow cytometry, and cross reactivity indexes – we performed principal component analysis (PCA). Interestingly, PCA projections of days 15 and 30 p.c. corroborate our assay-by-assay results. More specifically, both principal component projections separate most of the CD4+/DV2/CD4-/ZV and the double depleted groups (CD4-/DV2/CD4-/ZV) samples – most samples have positive values on PC2 on days 15 p.i. (Fig 10A and 10B). Also, most of the samples from the undepleted CD4+/DV2/CD4+/ZV group and from the CD4-/DV2/CD4+/ZV and CD4+/DV2/CD8-/ZV groups having in common the presence of CD4^+^ T cells before ZIKV infection cluster together – having positive values in PC1 and negative value in PC2 on day 15 p.c. and negative values in PC1 on day 30 p.c.–, which corroborates our results that demonstrates that these groups behave similarly at day 15 p.i.. Of particular interest is the group with CD4+ T cells depleted before primary DENV infection but not before ZIKV challenge. On day 15 p.c., half of samples (n=3) clustered very close with the double-depleted group with a positive value in PC1 similar to the other three samples from same group, but with a positive value in PC2 as the double depleted group.

**Fig 10.**
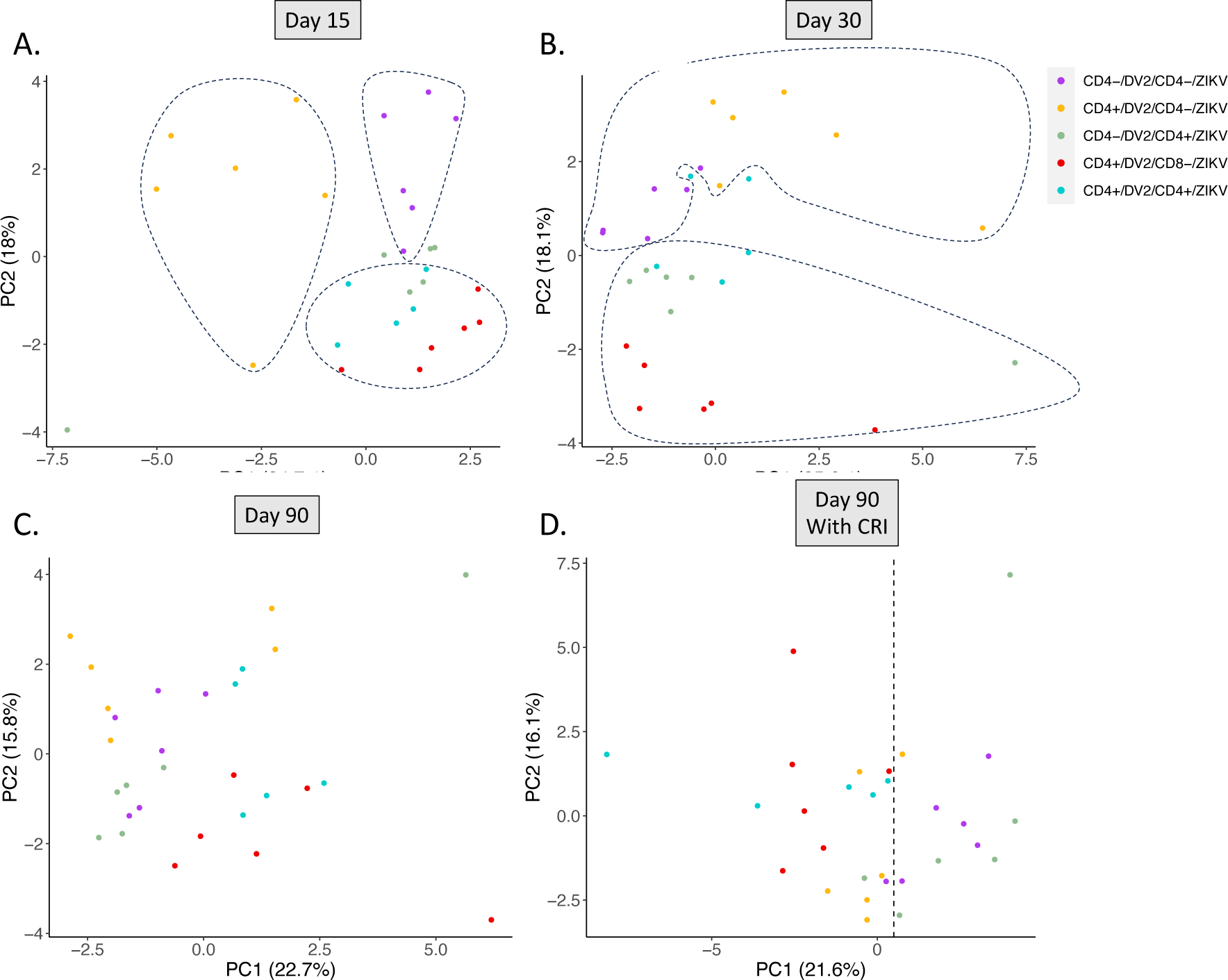
Principal component analysis of the effect of CD4+ T cell depletion on the ZIKV immune response. Principal component analysis was conducted in order to evaluate the overall effect of CD4+ T cells on various elements of the immune response against ZIKV including neutralization activity, IgM and IgG response and B and T cell response. Animal cohorts are depicted as follows based on treatment regime: CD4+/DV2/CD4+/ZV (shown in turquoise), CD4-/DV2/CD4+/ZV (shown in green), CD4+/DV2/CD4-/ZV (shown in yellow), CD4-/DV2/CD4-/ZV (shown in purple), and CD4+/DV2/CD8-/ZV (shown in red). Principal component analysis of day 15 **(A)**, day 30 **(B)** and day 90 **(C)** post ZIKV challenge. Principal component analysis of day 90 p.c. including CRI data **(D)**.

PCA projections of day 90 post ZIKV challenge shows how groups return to baseline, as there is no clear separation between groups (Fig 10C). This is in line with our individual assay-by-assay results that show that, after 90 days of infection, there is no significant difference between groups. Eighty three percent (83%) of the samples from the CD4-/DV2/CD4+/ZV showed negative PC1 clustering with 100% of samples from the double depleted group and 66% of samples from the group depleted of CD4+ T just before ZIKV infection (CD4+/DV2/CD4-/ZV). Nevertheless, after incorporating the CRI to our PCA matrix there is a dramatic shift with the CD4-/DV2/CD4+/ZV group (83% of samples) clustering together with all samples from the CD4-/DV2/CD4-/ZV double-depleted groups with most having now positive PC1 values of over 0.5 between most of the samples. This way those groups remained separated from the other groups (Fig 10D). These results are in line with our individual assay-by-assay results that show that, in most of the results the double depleted group and the CD4+/DV2/CD4-/ZV trend to have similar results while the CD4-/DV2/CD4+/ZV have similar profile from the two undepleted groups (CD4+/DV2/CD4+/ZV and CD4+/DV2/CD8-/ZV). Such a drastic change in the dimensionality of the data reflects the impact of the CRI and most importantly, this analysis strongly suggest a differential impact of CD4+ T cells on the Abs binding and neutralization properties versus specificity generating immune memory during a primary DENV infection.

Overall, PCA projections recapitulated the individual trends in the various analysis performed within our dataset. PCA further provides an excellent resource to corroborate the validity and biological importance of our results.

## Discussion

Despite many years or research, DENV pathogenies is not completely understood. However, it is recognized that a balanced humoral and T cell immune response is essential to drive protection.

The role of CD4^+^ T cells in DENV pathogenesis evolved over the years. Early data and subsequent works highlighted their role in DENV pathogenesis (Duangchinda et al., 2010; Mangada and Rothman, 2005; Rivino et al., 2013; Rouers et al., 2021; Yu et al., 2022). However, more recent works show a relationship between these cells and DENV protection (Graham et al., 2020; Weiskopf et al., 2015b). Also, a contribution of CD4^+^ T cells in protection against ZIKV has been documented (Cimini et al., 2017; Elong Ngono et al., 2019; Hassert et al., 2018; Lucas et al., 2018). Nevertheless, there are currently limited data available on the relationship between the generation of cross-reactive T cells following vaccination and protection from infection or severe DENV disease (Guy et al., 2008; Tricou et al., 2022; Waickman et al., 2019a) or ZIKV (Rivino and Lim, 2017; Wen and Shresta, 2017). In this work, we found that the presence of CD4^+^ T cells was essential for optimal control of primary DENV infection and an effective Abs switching, as the depletion of those cells resulted in increased mean viremia days and in a delay in anti-DENV IgM and IgG production. These results suggest a delay in B cells activation and in isotype class switching during primary DENV infection related to the lack of CD4^+^ T cells. Our findings are in contraposition with a prior work in mice showing that depletion of CD4^+^ T cells did not have a significant effect on viral clearance and were not required for the induction of DENV2-specific Abs or CD8+ T cell responses, and that the early anti-DENV2 Ab response is CD4^+^ T cells-independent (Yauch et al., 2010). Nevertheless, in agreement with our results, earlier works also performed in mice showed that depletion of CD4^+^ and CD8^+^ T lymphocytes significantly reduced survival of ACS46-primed mice challenged with the DENV JHA1 strain, and that Abs were not required to protect against DENV, suggesting that the cellular immune response targeting NS proteins may be a promising route in vaccine development against DENV (Amorim et al., 2016). In addition to their role interacting with B and CD8^+^ T cells, CD4^+^ T cells have also been shown to be protective against encephalitis induced by West Nile Virus (WNV), another flavivirus, in RAG(-/-) and C57BL/6 (H-2(b)) mice (Brien et al., 2008). Again, differences in animal models may account for the radical differences in our results from prior published works.

During secondary ZIKV challenge we identified a dynamic and sequential effect of prior DENV immunity and of CD4^+^ T cells as well. First, all groups showed a displacement of ZIKV peak viremia from day 1 and 2 p.c. to day 3 p.c. compared to naïve groups and second, we identified an early ZIKV viremia resolution in the CD4^+^ T cells undepleted group. This ZIKV viremia profile confirms an adverse effect of prior DENV immunity in ZIKV replication in comparison to DENV naïve animals as reported by our group before (Perez-Guzman et al., 2019; Serrano-Collazo et al., 2020). In addition to the preexisting DENV immunity, the depletions show a striking two-phase effect. Initially we identified an early significant ZIKV viremia control in the group having the higher significant CD4^+^ T cell frequency (as a compensatory response to the CD8^+^ T cell depletion), strongly suggesting an early role of DENV-experimented CD4^+^ T cells. However, during the mid-viremia period (days 5-7 p.c.) we identified a viremia rebound in this group with a strong trend to have higher AUC values compared to the groups having intact CD4^+^ and CD8^+^ T cell components. This change in profile, from an early significant ZIKV viremia control followed by a viremia rebound in this CD4^+^/CD8^-^ group, outlines a role for CD8^+^ T cells controlling viremia at later time points.

The result on the contribution of CD4^+^ T cells is coherent with prior reports confirming the role of those cells controlling ZIKV viremia in naïve but also in DENV-experimented animals (Wen et al., 2020). Mice depleted of CD4^+^T cells have more severe neurological sequela and significant increases in viral titers in the central nervous system. When those cells were transferred from ZIKV-immune mice, CD4+ T cells protected type I interferon receptor-deficient animals from a lethal challenge, showing that the response by these cells is necessary and sufficient for control of ZIKV disease (Hassert et al., 2018). On another work, depletion of CD4+ T cells in mice reduced plasma cell, GC B cell, and IgG responses to ZIKV without affecting CD8+ T cell response, and those cells were required to protect mice from a lethal dose of ZIKV after infection intravaginally, but not intravenously (Elong Ngono et al., 2019).

These results are also in line with our finding that depletion of CD4^+^ T cells before ZIKV challenge in animals with prior DENV immunity significantly dampened B cell immune response, delaying early IgM and IgG responses and ZIKV neutralization, supporting a protective role of CD4^+^ T cells against ZIKV mediated by different mechanisms.

Also interesting is our finding that depleting CD8^+^ T cells before ZIKV challenge resulted in a strong trend to have lower viremia, suggesting that CD8^+^ T cells do not play a crucial role in controlling early ZIKV infection. Prior several works support the role of CD8^+^ T cells and CR DENV virus-specific CD8^+^ T cells controlling ZIKV replication (Regla-Nava et al., 2018; Rivino and Lim, 2017; Wen et al., 2017; Wen and Shresta, 2017; Yauch et al., 2010). Our finding in the NHP model points out that DENV-experimented CD8^+^ T cells may have a role controlling ZIKV viremia but at later time points of infection. Our model suggests a sequential contribution of the cellular immune response with CD4^+^ and CD8^+^ T cells as early and late players controlling ZIKV viremia in DENV-immune individuals respectively.

Results from multiple CD4^+^ T cell depletions also shed light on the quantity and quality of the immune response in sequential DENV/ZIKV infections in the context of an optimal or suboptimal priming of B cells. It is known that CD4^+^ T cells can have a dissimilar contribution to the humoral immune response depending on the type of prior flavivirus exposure. A prior T cell receptor repertoire analysis revealed preferential expansion of CR clonotypes between JEV and ZIKV, suggesting that pre-existing immunity against JEV may prime the establishment of stronger CD4^+^ T cell responses to ZIKV purified-inactivated virus vaccination. These CD4^+^ T cell responses correlated with titers of ZIKV-nAbs in JEV-pre-vaccinated group, but not in flavivirus-naïve or YFV-pre-vaccinated individuals, suggesting a stronger contribution of CD4^+^ T cells in the generation of nAbs in the context of JEV-ZIKV cross-reactivity (Lima et al., 2021). This finding, together with our results and from others, emphasize that the efficacy of a DENV or ZIKV vaccine may also depend on the nature of prior or post exposure to flavivirus.

The binding or magnitude of neutralization against DENV2 or ZIKV after ZIKV challenge was essentially unaffected by the lack of CD4^+^ T cells before primary DENV infection. These results indicate that DENV-experimented CD4^+^ T cells do not contribute to an optimal humoral immune response against ZIKV and support prior reports confirming the specificity of ZIKV-induced humoral immune response in naïve (Collins et al., 2017; Duffy et al., 2009), but also in DENV-pre-exposed, subjects (Andrade et al., 2020). However, depletion before primary infection results in a significant increase in ZIKV-cross reactivity prior ZIKV challenge evidenced by the presence of CR ZIKV-binding IgG. This implies an essential role of CD4^+^ T cells to guarantee a more specific Abs repertoire after the *primo* infection. That role may contribute to avoid potential disease enhancement during a secondary flavivirus exposure, due to generation of wide CR DENV MBC as consequence of a suboptimal priming. In contrast to the total IgG, no animals had detectable anti-ZIKV-NS1 IgG Abs at baseline, indicating that those Abs are very specific, supporting the efforts to use this protein as a diagnostic marker (Castro-Trujillo et al., 2023; Guzman et al., 2010; Petphong et al., 2023; Vazquez et al., 1995), but also considering the antigen as a target for vaccination. However, removing CD4^+^ T cells prior to primary infection resulted in a significant increase in DENV anti-NS1 Abs durability. The implications of this finding need further considerations to determine their probable or not pathogenic role, during a subsequent natural exposure or DENV or ZIKV vaccine administration.

Despite this, NS1 has also been used as a strategy for protection against TBE (Beicht et al., 2023) and against JEV (Li et al., 2020; Nath et al., 2020; Zhou et al., 2020). Gonçalves et al. nicely showed in mice that the cooperation between CD4^+^ T cells and the humoral response induce a robust protection against DENV mediated by a DNA-NS1 vaccine (Goncalves et al., 2015). Rhesus macaques is a good model to evaluate the resppnse to NS1 protein as prior work using this specie and same DENV2 strain we used, confirmed that CD4^+^ T cells from these animals targeted NS1, NS3 and NS5 proteins (Mladinich et al., 2012).

Our results confirmed an enhanced circulation of DENV-NS1 antigen after DENV infection and significant lower DENV or ZIKV-specific anti-NS1 IgG response after the primary infection of challenge respectivelly, associated to the lack of CD4^+^ T cells, confriming a protective role for these cells.

Consistent with the Abs response, our data on the MBC and ACS after ZIKV challenge points to a *de novo* B cell response pathway that is T cell-dependent to promote the formation of both IgG MBCs and ASC subsets with good neutralizing capacity towards ZIKV. Only the groups with CD4^+^ T cells before the ZIKV challenge recorded a significant activation of both MBC and ACS, while the depleted groups showed a trend to activation but without reaching significant differences with their baseline frequencies. Previosly, we showed a similar profile (CD4^+^ T cells-dependent MBC and ACS activation) during tertiary flavivirus infection, but only in the sequence where the ZIKV challenge was preceded by a DENV infection and not when two consecutive DENV infections were secondary and terciary exposures (Marzan-Rivera et al., 2022). Our results are in agreetment with data from humans showing that ZIKV infection in DENV-naïve subjects (or with prior DENV immunity) induces both DENV-MBC and naive B cells with the production of ZIKV type-specific Abs in both cases (Andrade et al., 2020; Rogers et al., 2017). The recall memory against DENV2 detected in our work after the ZIKV challenge was signifincantly higher in the undpelted groups suggesting to be CD4^+^ T cells-dependent. However more specific studies on the BCR and Abs repertoir diversity, which are being imlemented by our group, are needed. A recent work characterized the plasmablast response in rhesus macaques exposed to primary ZIKV or sequential ZIKV/ZIKV or DENV3/ZIKV infections. Authors reported the highest VH and VL diversity levels upon primary ZIKV infection and lower levels in both homologous and heterologous secondary ZIKV or DENV responses (Singh et al., 2024). In addition to the authors discussion, those results also strongly suggest dissimilar T-B cells interactions in those different sequential flavirus infections.

A different scenario compatible with our prior findings on sequential DENV infections (Marzan-Rivera et al., 2022), showing that the pool of cross-reactive policlonal Abs were generated mainly from MBC-derived plasmablasts and not from naïve B cells, was characterized for repeated DENV infections (Xu et al., 2012).

Cell phenotyping results were consitent with the correlation analysis we performed. Only the groups that were CD4+ before ZIKV challenge showed a significant positive correlation between some pThf phenotypes and MBCs activation. However only the control undepeleted group showed significant correaltion with the activiation of the ACS. Interesting the group depleted of CD4^+^ T cells only before ZIKV infection showed a significan inverse correlation with the activation of the ACS. This finding and its relationship with the increased cross-reactivity in this group need further considerations. Interestignly, the group depleted of CD8^+^ T cells before ZIKV challenge – but with intact CD4^+^ T cells – did not showed any correlation with the activiation of these cells, suggesting a potential role of CD8^+^ T cells in an effective activation of the MBC and the ACS as it has been discussed before (Rouers et al., 2021). Finally we found that pTfh (CXCR5+) and pTfh with TH1 pehonotype (CXCR5+/CXCR3+), but not the the pTfh expressing the B cells markers activation (PD1+/ICOS+), had a strong correlation with EC50 values against ZIKV (but not DENV). However when we split the results by groups, only the control double positive group kept the significant correlation with the EC50 values, sugesting a contribution of the DENV-pre exposed T helper population in an effective neutralization against ZIKV. This also underscores a potential role for CD8^+^ T cells in quality of neutralization, a field largely unexplored in flavivirus research. In the context of DENV infection, one study showed how levels of CXCR3+CD8+ T cells had a significant positive correlation with levels of antigen specific MBCs at the post febrile phase after a DENV infection in adult patients in Singapore (Rouers et al., 2021).

Because CD4^+^ pTH cells are essential for the formation of germinal centers and to regulate germinal center B cell differentiation into plasma cells and memory B cells (Crotty, 2015), our correlation results support our other findings suggesting that the humoral immune response to ZIKV in DENV-experimented subject is most likelly T-cell dependent; any disruption or suboptimal T-B cells interactions during a primary DENV infection has consequence in the quality of the immune response to a secondary ZIKV exposure. Vaccination of naive HLA-DRB1*0101 transgenic mice with DENV/ZIKV-cross-reactive CD4^+^ T cell epitopes induced a CD4^+^ T cell response sufficient to reduce tissue viral burden following ZIKV infection suggesting that those cells expressing a TH1 pehnotype (as we characterized in our model) can suppress ZIKV replication, even in a Abs-independient manner (Wen et al., 2020). A recent work on the response to the TV003 vaccine in phase II clinical trial revealed that sero-positive subjects have increased baseline/pre-vaccination frequencies of circulating T follicular helper (cTfh) cells and, more important and inline we our findings, they found that the baseline/pre-vaccination cTfh profile correlated with the vaccines ability to launch neutralizing antibody response against all four DENV serotypes (Izmirly et al., 2022).

The findings on the correlation between the pTfh with the MBC and the magnitude of neutralization form our work have implications for vacines design. Lack of inducing an effective and specific CD4^+^ T cell (and most likely pTfh cells) responses might thus impair vaccine performance. Our results add on the mechanisms guiding the immune response during heterologous flavivrus infection and may contribute to rational vaccine development efforts.

One key finding from our work is the increased cross-reactivity to both DENV-EDIII domain and NS1 (but not to similar ZIKV regions) in absence of CD4^+^ T cells before the primary DENV infection but not before the ZIKV challenge. The fact that the cross reactivity was identified against both the structural EDIII domain and the NS1 protein agrees with the finding that CD4^+^ T cell epitopes in humans are skewed toward recognition of viral components that are also targeted by B lymphocytes and, from here, guiding the production of Abs mainly against the structural proteins (Rivino et al., 2013) or NS1 protein having also T and B cells epitopes (Aquino et al., 2023; Beicht et al., 2023; Perera et al., 2024; Shu et al., 2000). We do not completely understand why the cross-reactivity was higher against the NS1 protein than to the EDIII domain, but the factors responsible for directing CD4^+^ T cell to similar sites at the virus protein surface (or to NS1) in different flaviviruses are unknown (Aberle et al., 2018). However, they are most likely related to the structural context of these epitopes and their accessibility during antigen processing (Koblischke et al., 2017; Koblischke et al., 2018; Landry et al., 2017; Schwaiger et al., 2014). The involvement of NS1 can be explained as 50% or more of the MBC repertoire represented in mAbs from DENV patients bind to NS1, NS3, prM, or capsid in addition to the ENV (Beltramello et al., 2010; de Alwis et al., 2011). A prior work showed that half of anti-NS1 mAbs had limited cross reactivity among DENV, while most of the structural Abs showed full cross reactivity against all virus serotypes (Dejnirattisai et al., 2010). Supported in that data, the increased cross-reactivity against NS1 we found, reinforce the critical contribution of CD4^+^ T cells during a primary infection to the Abs specificity during a subsequent flavivirus exposure.

Strikingly, that increase in cross-reactivity was not accompanied by a detriment in Abs binding or DENV neutralization after the challenge, indicating that CD4^+^ T cells were not essential for those properties, but they were required for the Abs specificity. Interestingly, while CD8^+^ T cell depletion increased CD4^+^ T cell frequency, it had no effect on the specificity of epitope recognition, supporting that our finding is related to the quality of the priming. This suggest that the specificity of DENV Abs needs to be secured during a *primo* infection with an optimal B cell priming and may be regulated through a different mechanism than the binding and neutralization functions. After many years of extensive research, the mechanism regulating the Abs somatic hypermutation and affinity maturation are not well understood (Elsner and Shlomchik, 2020; Stebegg et al., 2018; Victora and Nussenzweig, 2012). How this process occurs in flavivirus is also largely unknown. However, we do know that high affinity Abs with higher SHM rate are generated in the GCs (Fink, 2012; Fink et al., 2007; Godoy-Lozano et al., 2016; Wong et al., 2020) where the encounter with the antigen-presenting cells, including CD4^+^ T cells) occurs. Our results agree and provide, for the first time, mechanistic evidence to prior findings where a population of broadly cross-neutralizing Abs in addition to the type specific neutralizing Abs was identified after secondary DENV infections (Patel et al., 2017).

Currently, our group is performing Ab depletion to characterize the TS vs. CR Abs and ADE experiment to test the potential enhancement of these cross-reactive Abs. We understand that results from these experiment will add instrumental information in support of our results. We also acknowledge that we used samples from day 90 p.c. where cross-reactive Abs start to decline but are still present (Katzelnick and Harris, 2017; Patel et al., 2017; Wahala and Silva, 2011). However, this will not change the fact that, in our experimental conditions, the Abs population after ZIKV challenge, in DENV-experimented individuals with a suboptimal T-B cell interaction, behave differently in their binding and neutralization properties vs. their specificity.

Also, experiments reproducing the sequence of infections ZIKV/DENV2 are underway, and exploring DENV/DENV and ZIKV followed by different DENV serotypes is in the pipeline.

Before, it has been highlighted that the risk of immunopathology associated with sub-optimal cross-reactive B-cell and T-cell responses to heterologous serotypes represent critical factors for the development of a fully protective DENV vaccine (Rivino and Lim, 2017). Our work adds on the relevance and uniqueness of the contribution of CD4^+^ T cells to the specificity of the neutralizing Abs to increase vaccines efficacy and to potentially prevent or decrease the risk of pathogenic manifestations mediated by ADE or similar mechanisms.

Finally, and not less relevant, our work reinforces the value of RMs as a reliable model with a strong translational capability and a needed complement to the results obtained with other animal models.

One limitation to advance our knowledge on the correlates of protection towards DENV is the lack of an adequate animal model to reproduce DENV clinical manifestations. Despite the proven value of the small animal models, most of the studies performing a fine dissection of the mechanisms addressing the protection against DENV and ZIKV use transgenic mice deficient in some essential anti-viral pathways (Yauch and Shresta, 2008; Zompi and Harris, 2012). The genetic modifications in murine models prevent to recognize the contribution of all components of the immune response as a whole and from here, the translation to humans may be narrow. In addition, it is known that the T cell immune response to flavivirus in humans is restricted by the HLA system (Angelo et al., 2017; Sierra et al., 2007; Weiskopf et al., 2016; Weiskopf et al., 2011; Wen et al., 2010; Zivna et al., 2002).

The value of rhesus macaues as NHPs to study the pathogenesis and immune response after natural infection, or vaccination, to DENV (Mladinich et al., 2012; Onlamoon et al., 2010; Osorio et al., 2014; Sariol et al., 2011; Sariol et al., 2007; Sariol and White, 2014; White et al., 2021; Young et al., 2023) or ZIKV (Abbink et al., 2017; Aliota et al., 2016; Best et al., 2017; Breitbach et al., 2019; Carroll et al., 2017; Coffey et al., 2018; Coffey et al., 2017; Dudley et al., 2019; Dudley et al., 2016; Haddow et al., 2017) is well documented. Also, our group and others have shown the usefulness of this model to dissect many aspects of DENV and ZIKV interactions (Crooks et al., 2021; George et al., 2017; Larocca et al., 2021; Marzan-Rivera et al., 2022; McCracken et al., 2017; Pantoja et al., 2017; Perez-Guzman et al., 2019; Serrano-Collazo et al., 2020; Valiant et al., 2018; Valiant et al., 2019). RMs have also shown to be very reliable to anticipate vaccine outcomes (Borges et al., 2019). Early immunization studies of RMs with Dengvaxia, either with the tetravalent formulation CYD-TDV (CYD1–4) or with two bivalent formulations (CYD1,2 + CYD3,4), resulted in higher responses against the dominant DENV4 and DENV1, and a very limited response to DENV2 serotypes in both approaches (Guy et al., 2009). Later on, results from humans showed a vaccine efficacy of 55.6%, 75.3% and 100.0% for serotypes 1, 3 and 4 respectively. The efficacy for serotype 2 was as low as 9.2 % resulting in an overall efficacy of 30.2% for all serotypes (Sabchareon et al., 2012). RM studies with DENVax (formulation used for TAK-003 vaccine) likewise anticipated the findings from the clinical trials. Different DENVax formulations failed to generate adequate responses against DENV4 and stimulated lower levels of nAbs to DENV 1, 3 and 4 but higher levels to DENV2 (Osorio et al., 2011a).

Overall, our findings have enormous impact for the planning and evaluation of DENV and ZIKV vaccine schedules, design, and monitoring, strongly suggesting the consideration of CD4+ T cell epitopes for the composition of new generation flavivirus vaccines.

## Acknowledgments

This work would have not been possible without the dedication and commitment of the Caribbean Primate Research Center and the Animal Resources Center staff. We thank Dr. Daniela Weiskopf for her valuable comments, Dr. Stephanie Dorta for her guidance in cell phenotyping and Dr. Espino for her support with the correlation analysis. The authors also recognize the excellent technical support provided by Dr. Nicole Marzán, Jorge Sánchez and Rafael Ocasio. Funding: This work was supported by R01AI148264 (NIAID) to C.A.S. and by P40 OD012217 to C.A.S. and M.I.M. (ORIP, OD, NIH), and partially by 5-U24-AI152170-04 to A.M.d.S. and R25GM061838 to A.M.-S.

## Author contributions

C.A.S., A.M.d.S, C.S.-C., A.M.-S. and L.C. developed the experimental design. C.R., M.I.M. and A.G.B. supervised and performed sample collection and animal monitoring. C.S.-C., A.M.-S., L.C., S.H. and L.A. performed the experiments. C.A.S., C.S.-C, A.M.-S., L.C. and M.S.-R. analyzed the data. C.A.S., C.S.-C, A.M.-S. and L.C drafted the article. C.A.S., C.S.-C., A.M.-S, L.C., S.H., M.S.-R., L.A., T.A., M.I.M., C.R., A.G.B. and A.M.d.S. reviewed and corrected the last version.

## Materials and Methods

### Experimental model

Young adult (4–7 years of age) male rhesus macaques (*Macaca mulatta*), seronegative for DENV and ZIKV, were housed in the CPRC facilities at the University of Puerto Rico, San Juan, Puerto Rico. All procedures were reviewed and approved by the Institute’s Animal Care and Use Committee at the Medical Sciences Campus, University of Puerto Rico (IACUC-UPR-MSC) and performed in a facility accredited by the Association for Assessment and Accreditation of Laboratory Animal Care (AAALAC) (Animal Welfare Assurance number A3421; protocol number, 7890116). In addition, the study was conducted following the USDA Animal Welfare Regulations, the Guide for the Care and Use of Laboratory Animals, and institutional policies, to ameliorate animal suffering by the recommendations of the Weatherall Report on the use of nonhuman primates in research. All procedures were conducted under anesthesia by intramuscular injection of ketamine at 10–20 mg/kg of body weight, as approved by the IACUC. Anesthesia was delivered in the caudal thigh using a 23-gauge sterile syringe needle. During the period of the entire study, the macaques were under the environmental enrichment program of the facility, also approved by the IACUC. Animals were continuously monitored by trained veterinarians at the Animal Research Center.

### Viral Stock

DENV-2 New Guinea 44 (NGC) strain (provided by Steve Whitehead, NIH/NIAID, Bethesda, MD) and ZIKV PRVABC59 (ATCC VR-1843) were used to infect macaques at different timepoints. Viruses were expanded and tittered by plaque assay and qRT-PCR using protocols standardized in our laboratories. In addition, both viruses were used for neutralization assays.

### Depletion treatment, viral infection, and sample collection

A total of 29 rhesus macaques seronegative for DENV and ZIKV were housed in the CPRC facilities, at the University of Puerto Rico, San Juan, Puerto Rico. During the first year, macaques were divided into two cohorts based on their depletion status. Twelve (12) animals were depleted of CD4+ T cells by administration of anti-CD4 Ab CD4R1 (NHP Reagent Resource; https://www.nhpreagents.org). Administration of the depletion treatment consisted of an initial s.c. inoculation of 50 mg/kg at 15 days prior to viral infection, followed by two i.v. injections of 7.5 mg/kg 12 days and 9 days before viral infection respectively. The seventeen (17) remaining animals received phosphate-buffered saline (PBS) as a control. Both cohorts were then infected with DENV-2 NGC strain (5.0 x 10^5^ pfu s.c.), which is denoted from here on as the primary infection. Following one year after infection, animals were further divided into five cohorts based on depletion status. Animals were depleted of CD4+ or CD8+ T cells, or treated with PBS, followed by a secondary infection with ZIKV (1 × 10^6^ pfu s.c.). CD8+ T cell depletion was performed by administration of anti-CD8 alpha MT807R1 at the same dosage as CD4+ T cells depletion treatment. CD4+ T cell depletion was performed on groups 1 and 2 (group 1: n=6, denoted CD4-/DV2/CD4-/ZV; group 2: n=6, denoted CD4+/DV2/CD4-/ZV). Group 3 was depleted of CD8+ T cells (n=6, denoted CD4+/DV2/CD8-/ZV). Lastly, the remaining two groups were administered PBS (group 4: n=6, denoted CD4-/DV2/CD4+/ZV; group 5: n=5, denoted CD4+/DV2/CD4+/ZV).

Blood samples were collected on days ^-^15, ^-^12 and ^-^9 pre-infection, and during the first fifteen consecutive days after infection, followed by a monthly sample collection from there onwards until a year later when sample collection pattern described is repeated for the secondary infection. Collected samples included serum, heparinized whole blood, and citrate whole blood for PBMCs isolation. Macaques were monitored after treatments and infections by trained veterinarians for evidence of disease and clinical status. Rectal and external temperatures and weights were taken on day 0 and every other day pre- and post-infection. A summary for the experimental design and groups is provided (SFig 1).

### Immunophenotyping

Phenotypic characterization of Rhesus macaque’s adaptive immune response was performed by 16-multicolor flow cytometry using fluorochrome-conjugated Abs at several time points. Depletion efficacy was confirmed by measuring the frequency of CD4^+^, CD8^+^ T cells, and CD20^+^ B cells on the following timepoints: pre-treatment/depletion baseline, and days 2, 5, and 9 post-treatment. In addition, B and T cell frequencies were measured on baseline and days 1, 2, 3, 7, 15, 30, 60 and 90 p.i. and p.c.. Aliquots of 150 uL of heparinized whole blood were incubated with a mix of antibodies for 30 min. in the dark at room temperature. After incubation, red blood cells are fixed and lysed with BD FACS fix and lyse solution, and cells were washed twice with BSA 0.05%. Samples were analyzed using a MACSQuant^®^ Analyzer 16 Flow Cytometer (Miltenyi Biotec, CA). Antibodies used in this study were: CD8-FITC (DK25) from Sigma; CD3-PerCP (SP34), Ki67-Viogreen (B56), CD4-APC (L200), CD69-PeCy7 (FN50), CD95-PE (DX2), IgG-APC-H7 (G18-145), CD11b-AF 700 (ICRF44), CD80-BV650 (L307.4), CD27-PE (M-T466) from BD Biosciences; CD20-PacBlue (2H7), HLA-DR-BV570 (L243), CD138-FITC (DL-101), ICOS-PE/Dazzle 594 (C398.4H), CXCR3-PECy7 (GO25H7), PD-1-BV650 (EH12.2H7) from Biolegend; CD28-APC-Vio770 (I5E8) and CCR7-PerCP Vio770 (REA546) from Miltenyi; CXCR5-PE (MU5UBEE) from Thermo eBiosciences; and CD38-APC (AT-1) from Adipogen.

For analysis, lymphocytes (LYM) were gated based on their characteristic forward and side scatter patterns. B cells were defined as CD20^+^CD3^-^, Memory (MBC= CD20^+^, CD3^-^, CD27^+^), class-switched IgG Memory (IgG MBC= CD20^+^, CD3^-^, CD27^+^, surface IgG+ (sIgG), plasmablasts as (PB= CD20^-^/^low^, CD3^-^, CD11b^-^, HLA-DR^+^, CD80^+^) as described by Silveira et al., 2015 (Silveira et al., 2015), plasma cells as (PC= CD20^-^/^low^, CD3^-^, CD138^+^) as described by Martinez-Mirullo et al., 2016 (Martinez-Murillo et al., 2016). Gating strategy for B cells and subsets is presented on Supplementary Figure 9 (SFig 9). T cells were defined as CD3^+^CD20^-^, CD4^+^ T cells (CD3^+^CD20^-^CD4^+^), CD8^+^ T cells (CD3^+^CD20^-^CD8^+^), and subpopulations were defined inside the CD3^+^ T cells. Naïve (CD3^+^CD20^-^ CD28^+^CD95^-^), EM (CD3^+^CD20^-^CD28^-^CD95^+^), and CM (CD3^+^CD20^-^CD28^+^CD95^+^) T cell subpopulations were determined within CD4^+^ and CD8^+^ T cells. Gating strategy for T cells and memory subsets is presented on Supplementary Figure 10 (SFig 10), while gating strategy for the characterization of CD4^+^ pTfh cells was represented by selecting CM and EM T cells (CD4^+^/CXCR5^+^/CXCR3^+^) expressing activation and migration markers such as (CCR7^+^/PD-1^+^/ICOS^+^) (SFig 11). All data from cell population frequencies and immune markers expression were analyzed using Flow/Jo software and GraphPad Prism (FlowJo LLC Ashland, OR).

### qRT-PCR

DENV-2 and ZIKV viral RNA for real-time PCR assay was extracted from 200uL of virus isolate (previously tittered by focus reduction neutralization assay) and from acute serum samples using a MagMax Viral/Pathogen Nucleic Acid Isolation Kit (Applied Biosystems, MA) as per the manufacturer’s instructions. Viral RNA extraction was performed using a KingFisher Duo Prime Purification System (Thermo Fisher Scientific, MA). In brief, a Viral Pathogen DNA/RNA binding magnetic bead solution is prepared, 200uL of sera, 500uL of Viral Pathogen Nucleic Acid isolation kit wash buffer, 5uL of a protein kinase solution, and 1mL of ethanol solution (80% ethanol in DNAse free water) were added to specified wells in a KF Duo Deepwell 96 plate (Thermo Fisher Scientific, MA) and loaded on the KingFisher Duo Prime Purification System. Purified RNA was collected and stored at -80°C in kit elution buffer solution. Real-time PCR for DENV-2 and ZIKV isolated RNA was carried out using the Gensig Real-Time PCR detection kit for Dengue Virus and the Gensig Real-Time PCR detection kit for Zika Virus (Primer Design, UK), respectively, as per the manufacturer’s instructions. Briefly, a kit Positive Control is diluted in 500uL of a kit Template buffer and a series of tenfold dilutions were prepared as follows: 2 x 10^5^/uL, 2 x 10^4^/uL, 2 x 10^3^/uL, 2 x 10^2^/uL, 20/uL, 2/uL (copy number/ mL). For the reaction mix, 10uL of oasig 2X Precision one step qRT-PCR was combined with 1uL of kit probe/primer mix and 4uL of sterile water, for a total volume of 20uL once the RNA sample was added. The reaction mix was added to a 96 well plate, alongside the experimental RNA samples, a kit non-template control (NTC), an RNA extraction negative control (DNAse free water), a positive control (RNA extraction from virus stock), and calibration standards, in specified wells. The plate was then loaded, and the RT-PCR reaction was performed using a Quant Studio 5 Real-Time PCR Instrument (Applied Biosystems, MA). For quantification, a standard curve was generated from the ten-fold dilutions of the kit RNA positive control.

### ELISAs for DENV and ZIKV

Serostatus of animals for DENV and ZIKV was assessed before and after DENV-2 infection using DENV IgG/IgM commercial kits (Focus Diagnostics, CA). ZIKV serostatus was also assessed using a commercial kit for ZIKV IgM (InBios, International Inc, Seattle) and ZIKV IgG (XpressBio, Frederick, MD). Timepoints measured include baseline and days 7, 10, 15, 30, 60, and 90 post-infection. All assays were performed per the manufacturer’s instructions and as described by our group (Pantoja et al., 2017; Perez-Guzman et al., 2019; Serrano-Collazo et al., 2020).

### ELISAs for detection of DENV NS1 antigen, DENV anti-NS1 IgG and ZIKV anti-NS1 IgG

Detection of DENV NS1 antigen after primary DENV-2 infection, ZIKV anti-NS1 IgG after secondary ZIKV infection, and DENV anti-NS1 IgG after both primary and secondary infections were measured using commercial kits (InBios, International Inc, Seattle). The assays were performed per the manufacturer’s instructions. Timepoints measured include baseline and days 3, 5, 7, 15 and 30 p.i. for DENV NS1, and baseline and days 15, 30 and 90 p.c. for ZIKV and DENV anti NS1 IgG.

### DENV neutralization assays

For the Focus Reduction Neutralization Test (FRNT) for DENV, Vero81 cells (ATCC CCL-81) were seeded at approx. 2.0 x 10^5^ per well in 24 well-plates with DMEM (Dulbecco’s Modified Eagle’s medium, Thermo Fisher Scientific) for approx. 18 hours. Serum dilutions (ten-fold) were prepared in diluent medium Opti-MEM (Invitrogen) with 2% FBS (Gibco) and 1% antibiotic/antimycotic (Hyclone). Virus was added to each dilution and incubated for 1 hour at 37°C/5%. Before inoculation, the growth medium was removed, and cells were inoculated with 50 uL per well of serum-virus dilution in triplicates; plates were incubated for 1 hour at 37°C/5%CO_2_/rocking. After incubation, 125 uL per well of overlay (Opti-MEM 1% carboxymethylcellulose (Sigma), 2% FBS, 1% non-essential amino acids (Gibco) and 1% antibiotic/antimycotic (HyClone)) was added to the plates containing virus dilutions, followed by an incubation period of 48 hours at 37°C/5%CO_2_. After two days, overlay was removed with PBS and fixed with 4% paraformaldehyde for 30 minutes. Plates were blocked with 5% non-fat dairy milk in 1X perm buffer (BD Cytofix/Cytoperm) for 10 min and incubated for 1hr/37°C/5%CO2/rocking with anti-E mAb 4G2 and anti-prM mAb 2H2 (kindly provided by Dr. Aravinda de Silva and Ralph Baric, University of North Carolina Chapel Hill, NC, USA), both diluted 1:100 in blocking buffer. Plates were washed three times with PBS and incubated 1hr/37°C/5%CO_2_/rocking with horseradish peroxidase (HRP)-conjugated goat anti-mouse antibody (Sigma), diluted 1:1,500 in blocking buffer. Foci were developed with TrueBlue HRP substrate (KPL) and counted using an Elispot reader. Results were reported as FRNT with a 60% or greater reduction in DENV foci (FRNT60). A positive neutralization titer was designated as 1:20 or greater, while ˂1:20 was considered a negative neutralization titer.

### ZIKV neutralization assays

For the Plaque Reduction Neutralization Test (PRNT) for ZIKV, Vero81 cells (ATCC CCL-81) were seeded at approx. 2.0 x 10^5^ per well in 24 well-plates with DMEM (Dulbecco’s Modified Eagle’s medium, Thermo Fisher Scientific) for approx. 18 hours. Serum dilutions (ten-fold) were prepared in a diluent medium (Opti-MEM (Invitrogen) with 2% FBS (Gibco) and 1% antibiotic/antimycotic (Hyclone). Virus was added to each dilution and incubated for 1 hour at 37°C/5%/CO2. Before inoculation, the growth medium was removed, and cells were inoculated with 100 uL per well of each dilution in triplicates; plates were incubated for 1 hour at 37°C/5%CO2/rocking. After incubation, 1mL per well of overlay (Opti-MEM 1% carboxymethylcellulose (Sigma), 2% FBS, 1% non-essential amino acids (Gibco), and 1% antibiotic/antimycotic (HyClone)) was added to the plates containing virus dilutions, followed by an incubation period of 4 days at 37°C/5% CO2. After four days of incubation at 37°C/5%CO2, the overlay was removed; the cells were washed twice with phosphate-buffered saline (PBS), fixed in 80% methanol, and stained with crystal violet and foci were counted. The mean focus diameter was calculated from approx. twenty foci per clone were measured at 35 magnifications. Results were reported as the PRNT with a 60% or greater reduction in ZIKV or DENV plaques (PRNT60). A positive neutralization titer was designated as 1:20 or greater, while ˂ 1:20 was considered a negative neutralization titer.

### EDIII and NS1 Antigen Production

ZIKV (H/PF/2013) EDIII antigen used in this study was produced as previously described (Adams et al., 2021). EDIIIs of DENVs were also cloned and expressed similarly to ZIKV EDIII. Briefly, codon-optimized genes encoding the EDIIIs of DENV1 (AAB70694·1, E protein aa 297-394), DENV2 (ADA00411·1, E protein aa 297-394), DENV3 (AAB69126·2, E protein aa 295-392) and DENV4 (ADA00410·1, E protein aa 297-394) with an N-terminal human serum albumin signal peptide, a polyhistidine tag, and a HaloTag were cloned into the mammalian expression plasmid pαH. Recombinant DENV EDIII antigens were expressed in mammalian Expi293 cells and purified from the culture supernatant using Ni-NTA agarose (Qiagen). Purified EDIII antigens were biotinylated using HaloTag PEG biotin ligand (Promega), according to the manufacturer’s protocol. For NS1 antigen production, codon-optimized genes encoding the NS1s of DV1 NS1 Region 37, DV2 NS1 Region 39, DV3 NS1 Region22, DV4 NS1 Region 45 and ZKVNS1 Region 47 were used. Mobility-shift analysis was used to assess the biotinylated EDIII antigens using SDS-PAGE. The mammalian expression plasmids and sequences will be made available in the plasmid repository and Genbank, respectively.

### Coupling antigens to beads via avidin-biotin interaction

Site-specifically biotinylated EDIII antigens were coupled to unique MagPlex®-Avidin Microspheres (DENV1 (region 12), DENV2 (region 26), DENV3 (region 36), DENV-4 (region 38) and ZIKV (region 30)) at a concentration of 5 μg of antigen per 106 beads in phosphate-buffered saline, pH 7.4 (PBS, 137 mM NaCl, 2.7 mM KCl, 8 mM Na2HPO4, and 2 mM KH2PO4) for 1 hour at 37°C with shaking at 700 rpm. Beads were then washed/blocked three times in block buffer (PBS + 1% BSA) and resuspended in wash buffer at 2×106 beads/mL.

### EDIII and NS1 multiplex Assay

DENV and ZIKV EDIII coupled beads were combined in equal ratios by plating 50 μL containing 2,500 beads per antigen into each well of a 96-well assay plate (Cat. No:655906). Human serum samples were diluted 1:500 in block buffer. Mouse monoclonal antibodies specific for each EDIII antigen (ZV-67 (ZIKV), E103 (DENV1), 3H5 (DENV2, 8A1 (DENV3), E88 (DENV4)) and NS1 antigen were titrated in each assay as controls for assay consistency across couplings, plates, and assay days. A magnetic plate separator was used to remove the block buffer from the assay plate, and 50 μL of diluted serum or mAb was added to the beads. The serum and bead mixture were incubated for 1 hour at 37°C with shaking at 700 rpm. Beads were washed thrice200 μL block buffer per well using a magnetic separator. 50 μl of Goat anti-Human IgG Fc Multi-species SP ads-PE (Southern Biotech, catalog:2014-09) and Goat anti-Mouse IgG Fc Human ads-PE (Southern Biotech, catalog:1030-09) were added to the beads (6 μg/ml in block buffer) and incubated for 1 hour at 37°C with shaking at 700 rpm to detect respective antibody species. Beads were washed thrice with 200 μL block buffer per well using a magnetic separator and resuspended in 100 μL of block buffer before acquiring data. Fluorescence data were obtained using a Luminex 200 analyzer set to acquire at least 50 beads per bead region.

### Principal Co1mponent Analysis (PCA)

All PCA analyses were performed using R statistical software R version 4.2.3 using the *prcomp* function of the *stats* package (Team et al., 2018; Team, 2010). To examine global patterns of infection, we carried out a PCA using all the parameters measured during the infection timeline; these analyses were performed for day 15, day 30 and day 90 of infection. The parameters used to perform PCA are depicted in Supplementary Table 3 (Table S3).

### Quantification and statistical analysis

Statistical analyses were performed using GraphPad Prism 9.0 software (GraphPad Software, San Diego, CA, USA). For viral burden analysis, the log titers and levels of vRNA were analyzed by multiple unpaired t-tests and two-way ANOVA. The statistical significance between or within groups evaluated at different time points was determined using one-way or two-way analysis of variance (ANOVA) (Tukey’s, Sidak’s, or Dunnett’s multiple comparisons test) or unpaired t-test to compare the means. Significant multiplicity-adjusted p-values (*< 0.05, **< 0.01, ***< 0.001, ****< 0.0001) show statistically significant differences between groups (Tukey test) or time points within a group (Dunnett test).

## SUPPLEMENTARY FIGURES

**SFig 1.**
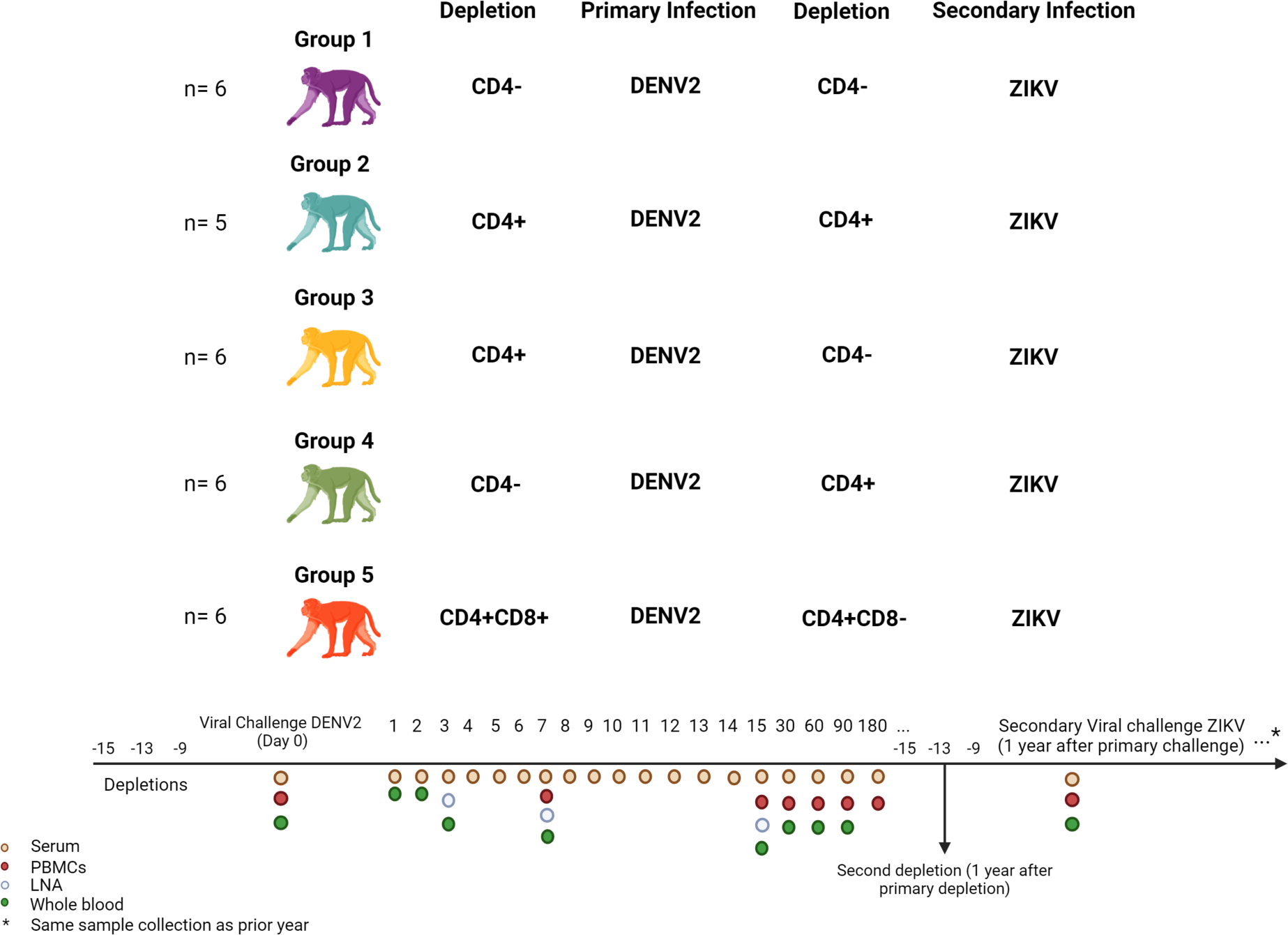
Experimental design of T cell depletion and heterologous DENV2 and ZIKV infections in flavivirus-naïve macaques. Five cohorts of rhesus macaques (*Macaca mulatta*) underwent depletion of CD4+ T cells (or PBS in control group) followed by DENV-2 (5 x 10^5^ pfu s.c.) infection at different timepoints. One year later, all cohorts were exposed to ZIKV strain PRABCV59 (1 x 10^6^ pfu s.c.) after a second round of CD4+ or CD8+ T cell depletion (or PBS in control group). Sample collection was performed at various time points up to 90 days post-infection (p.i.) for serum, whole heparinized blood, and PBMC isolation.

**SFig 2.**
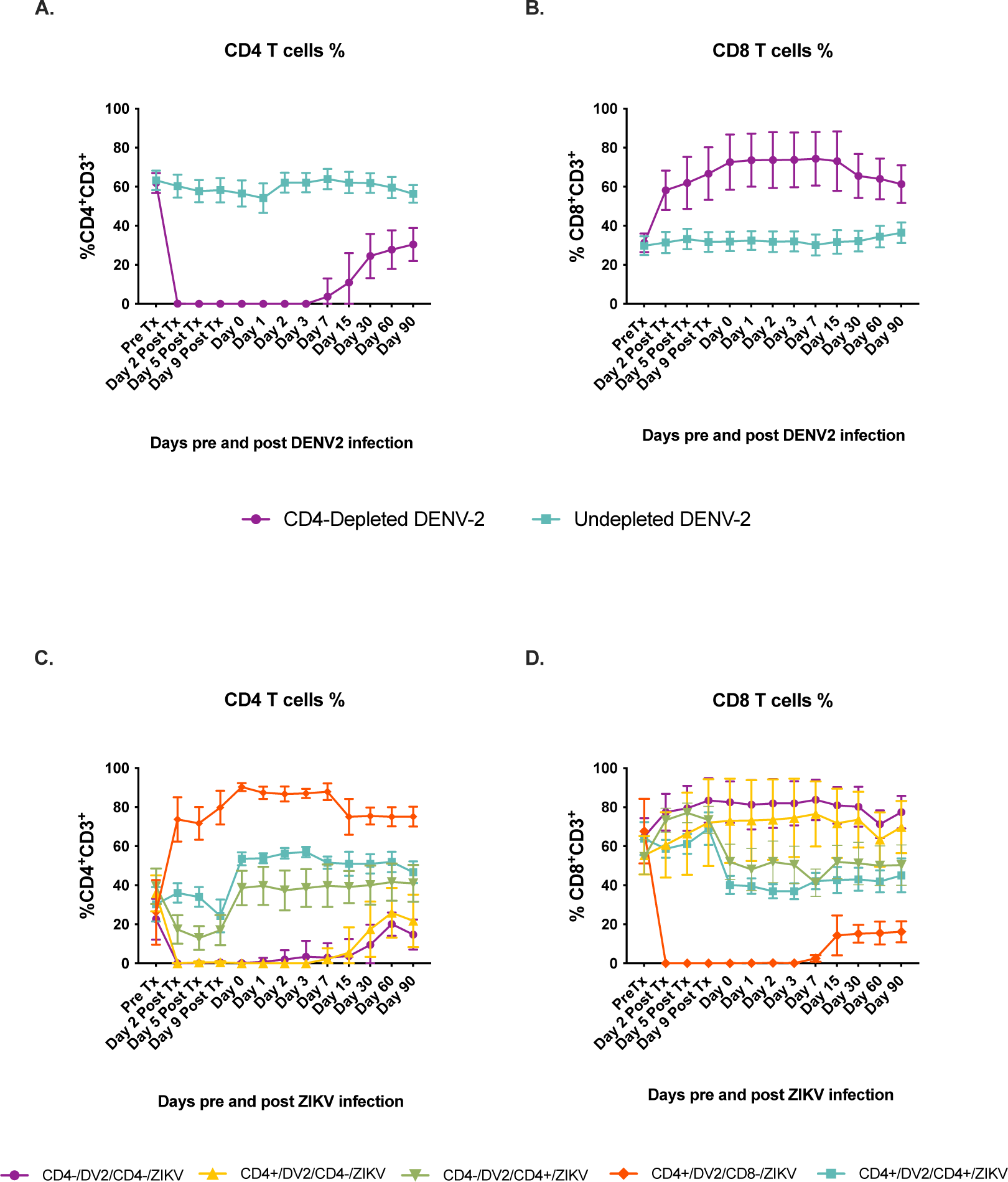
CD4+ and CD8+ T cell frequency following depletion treatments prior to DENV2 and ZIKV infections. Frequency of CD4+ and CD8+ T cells was assessed by immunophenotyping before and after depletion treatments using flow cytometry. **Upper panel**. CD4-depleted animals are depicted in purple and undepleted animals are depicted in turquoise. CD4+ **(A)** and CD8+ **(B)** T cell depletion efficacy for depleted and undepleted animals before, during and after DENV2 infection. **Lower panel**. Animal cohorts are depicted as follows based on treatment regime: CD4+/DV2/CD4+/ZV (shown in turquoise), CD4-/DV2/CD4+/ZV (shown in green), CD4+/DV2/CD4-/ZV (shown in yellow), CD4-/DV2/CD4-/ZV (shown in purple), and CD4+/DV2/CD8-/ZV (shown in red). CD4+ **(C)** and CD8+ **(D)** T cell depletion efficacy for depleted and undepleted animals before, during and after ZIKV infection.

**Table S1.**
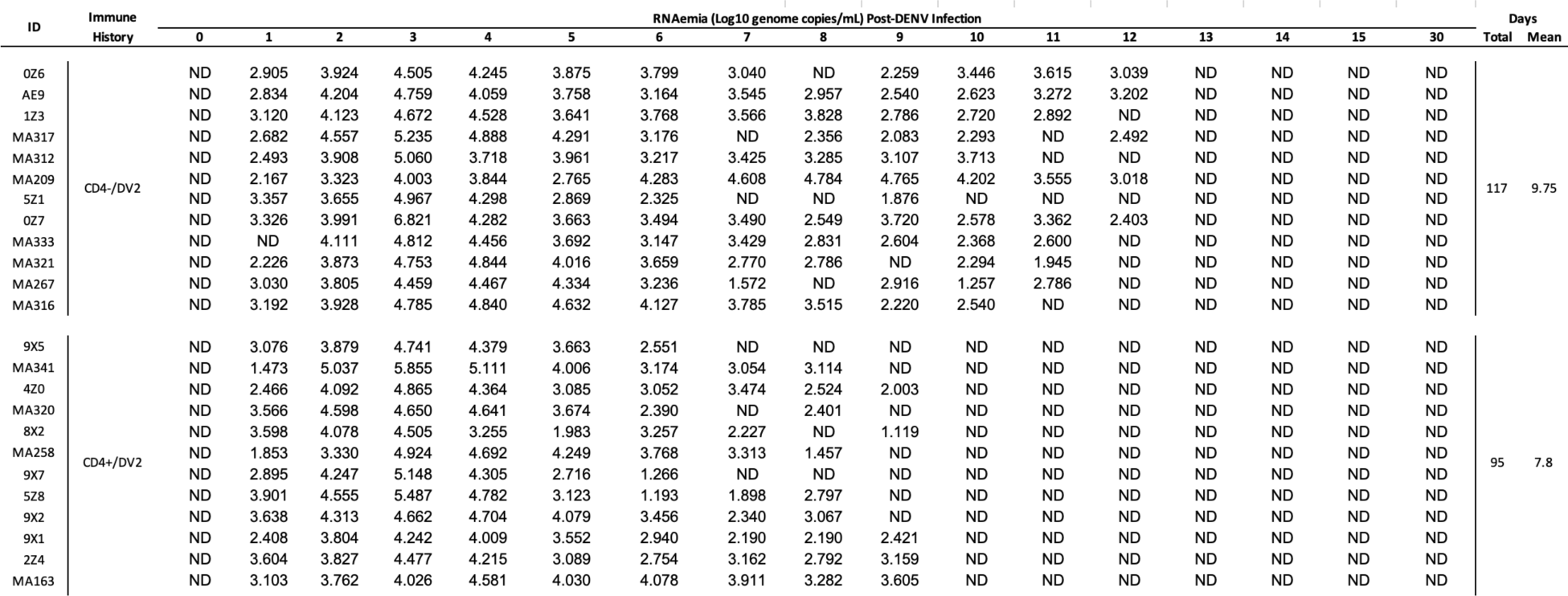
DENV-2 RNAemia of CD4-depleted and undepleted animals. DENV-2 RNA detection during the first 15 days and day 30 p.i. Mean viremia days per group was calculated using days with detectable RNAemia divided by the number of animals in each group. ND= viral RNA not detected. Blank spaces represent sample was able to be collected.

**Table S2.**
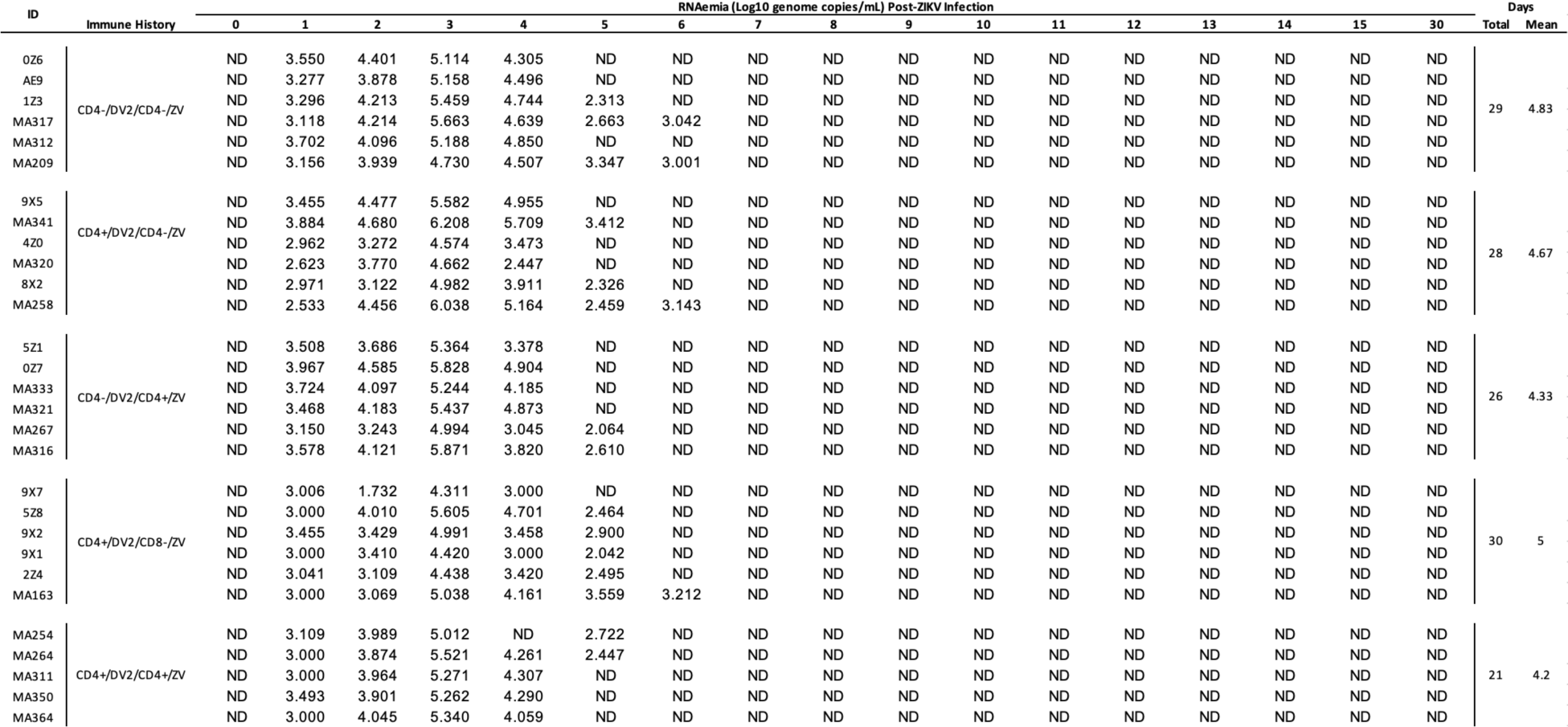
ZIKV RNAemia days of DENV-immune macaques with different T cell depletion status. ZIKV RNA detection during the first 15 days and day 30 p.i. Mean viremia days per group was calculated using days with detectable RNAemia divided by the number of animals in each group. ND= viral RNA not detected.

**SFig 3.**
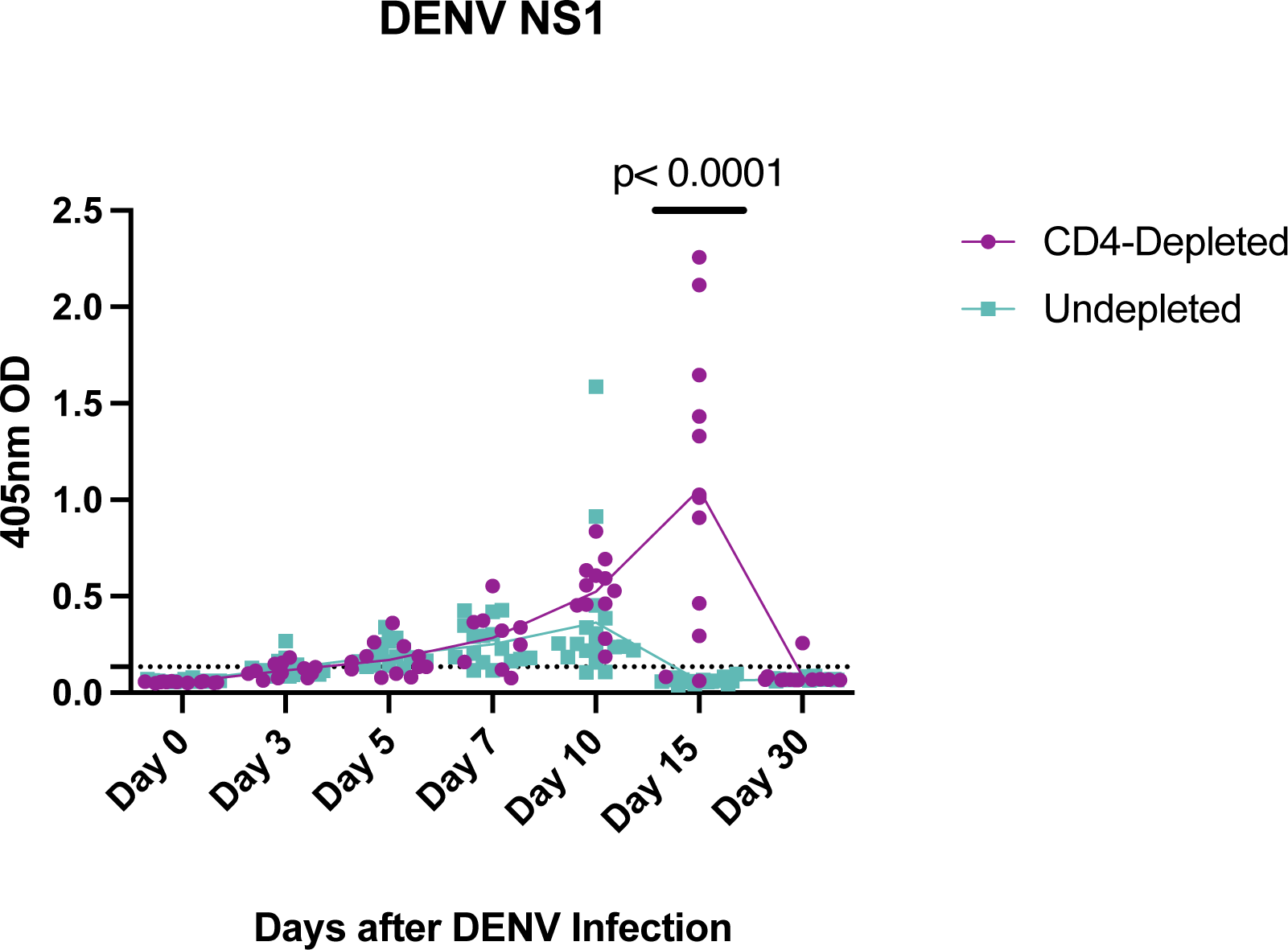
DENV NS1 levels are affected by absence of CD4+ T cells after DENV-2 infection. DENV-NS1 levels after DENV-2 infection were measured using a commercial ELISA test. CD4-depleted animals are depicted in purple and undepleted animals are depicted in turquoise. Dotted lines indicate the limit of detection for each test. Statistically significant differences among and within groups were calculated by two-way ANOVA using Tukey’s multiple comparisons test and unpaired t-tests.

**SFig 4.**
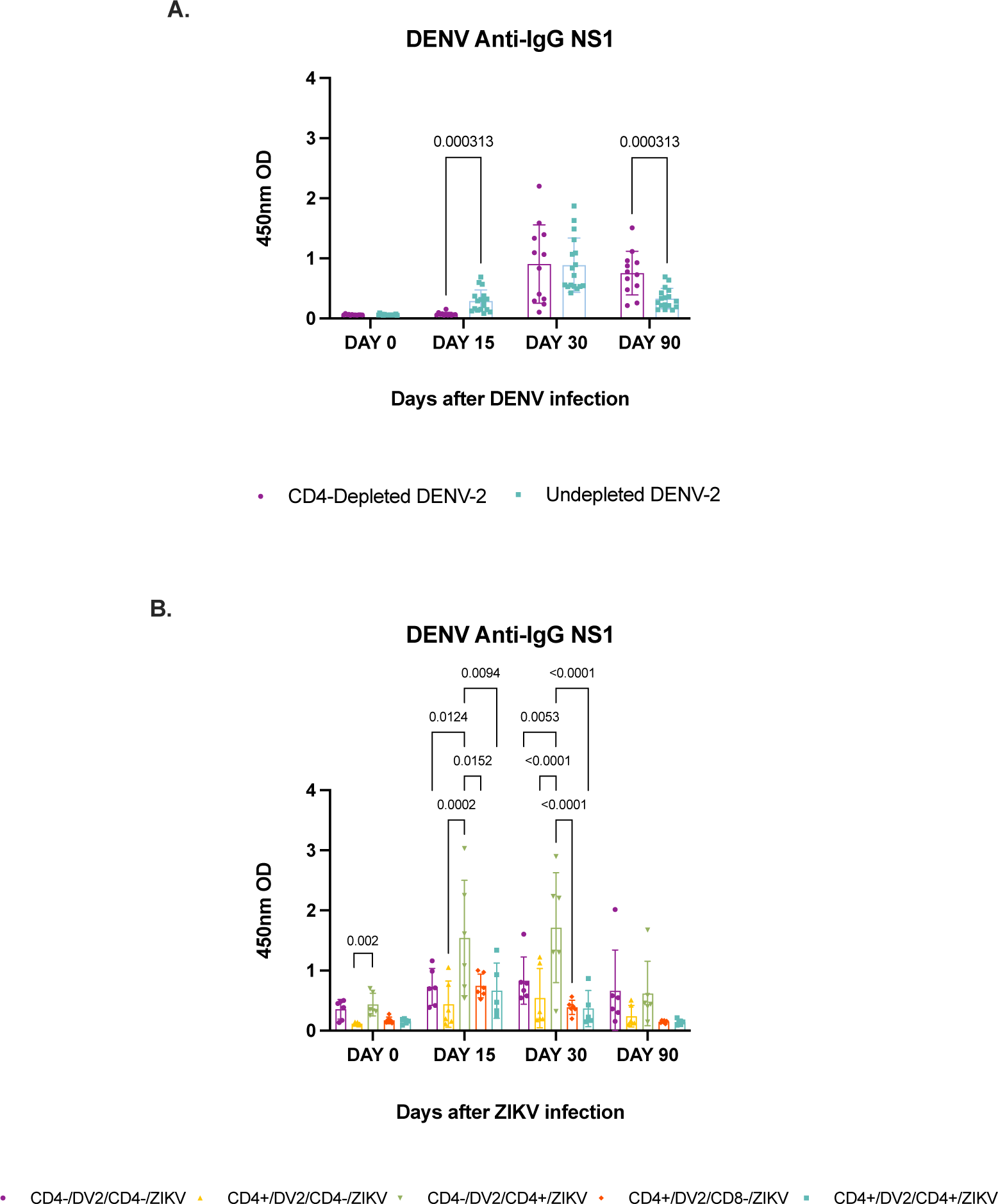
Depletion of CD4+ T cells before DENV infection increases DENV NS1 anti-IgG levels. DENV-NS1 anti-IgG levels were measured using commercial ELISA tests. **(A)** CD4-depleted animals are depicted in purple and undepleted animals are depicted in turquoise. DENV NS1 anti-IgG levels after DENV-2 infection. **(B)** Animal cohorts are depicted as follows based on treatment regime: CD4+/DV2/CD4+/ZV (shown in turquoise), CD4-/DV2/CD4+/ZV (shown in green), CD4+/DV2/CD4-/ZV (shown in yellow), CD4-/DV2/CD4-/ZV (shown in purple), and CD4+/DV2/CD8-/ZV (shown in red). DENV NS1 anti-IgG levels after ZIKV infection. No limit of detection or cut-off value is provided because this value will vary depending on flavivirus disease prevalence in the geographical location where the test is performed. Statistically significant differences among and within groups were calculated by two-way ANOVA using Tukey’s multiple comparisons test and multiple t-tests.

**SFig 5.**
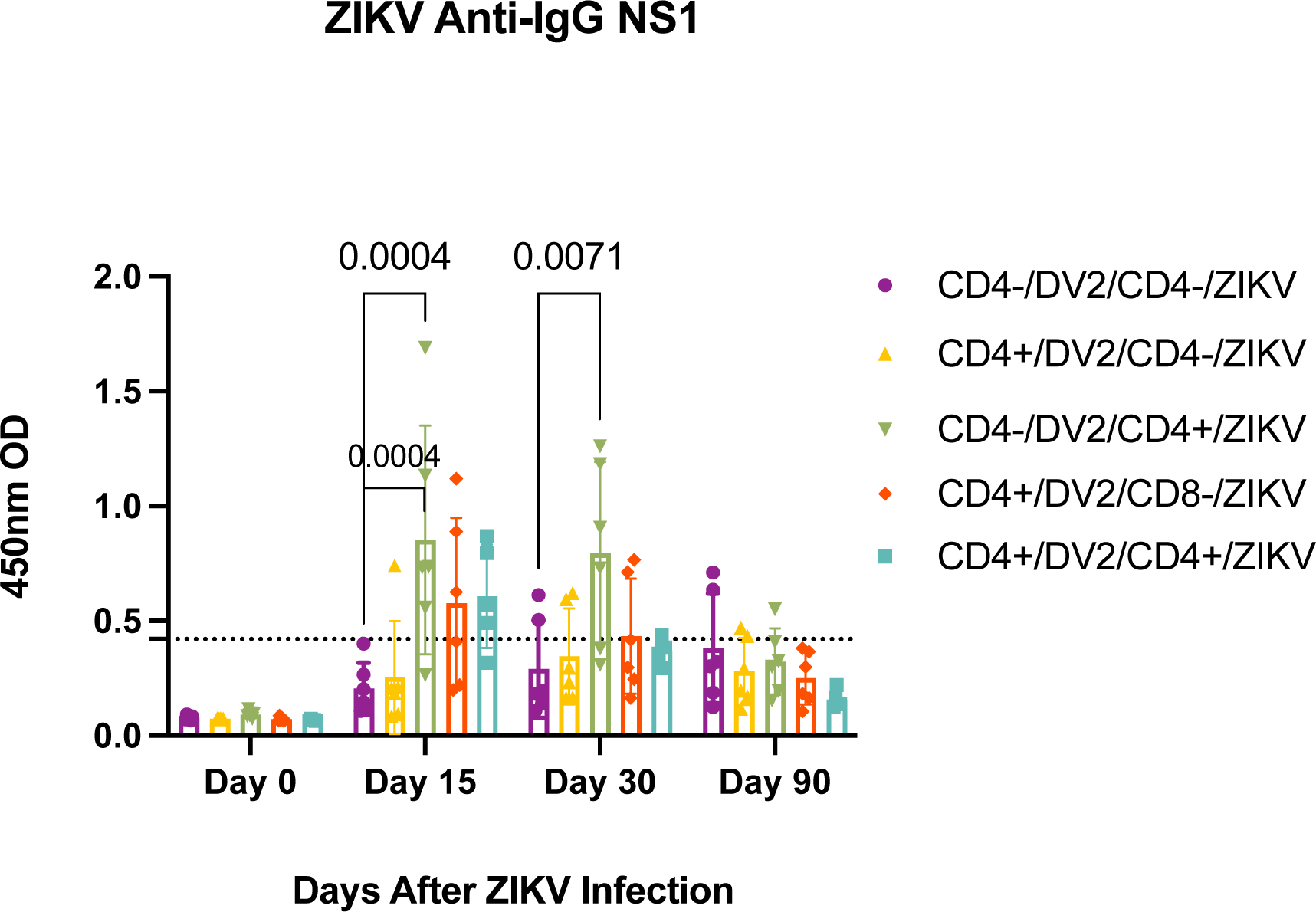
CD4+ T cell depletion status prior to DENV infection has no impact in production of ZIKV NS1-binding IgG after ZIKV infection. ZIKV NS1-IgG response after ZIKV infection in the presence of DENV immunity was assessed using a commercial ELISA test. Animal cohorts are depicted as follows based on treatment regime: Animal cohorts are depicted as follows based on treatment regime: CD4+/DV2/CD4+/ZV (shown in turquoise), CD4-/DV2/CD4+/ZV (shown in green), CD4+/DV2/CD4-/ZV (shown in yellow), CD4-/DV2/CD4-/ZV (shown in purple), and CD4+/DV2/CD8-/ZV (shown in red). Dotted lines indicate the limit of detection for each test. Binding capacity of ZIKV NS1-IgG antibodies from CD4-depleted and undepleted animals after ZIKV infection. Statistically significant differences among and within groups were calculated by two-way ANOVA using Tukey’s multiple comparisons test and unpaired t-tests.

**Sfig 6.**
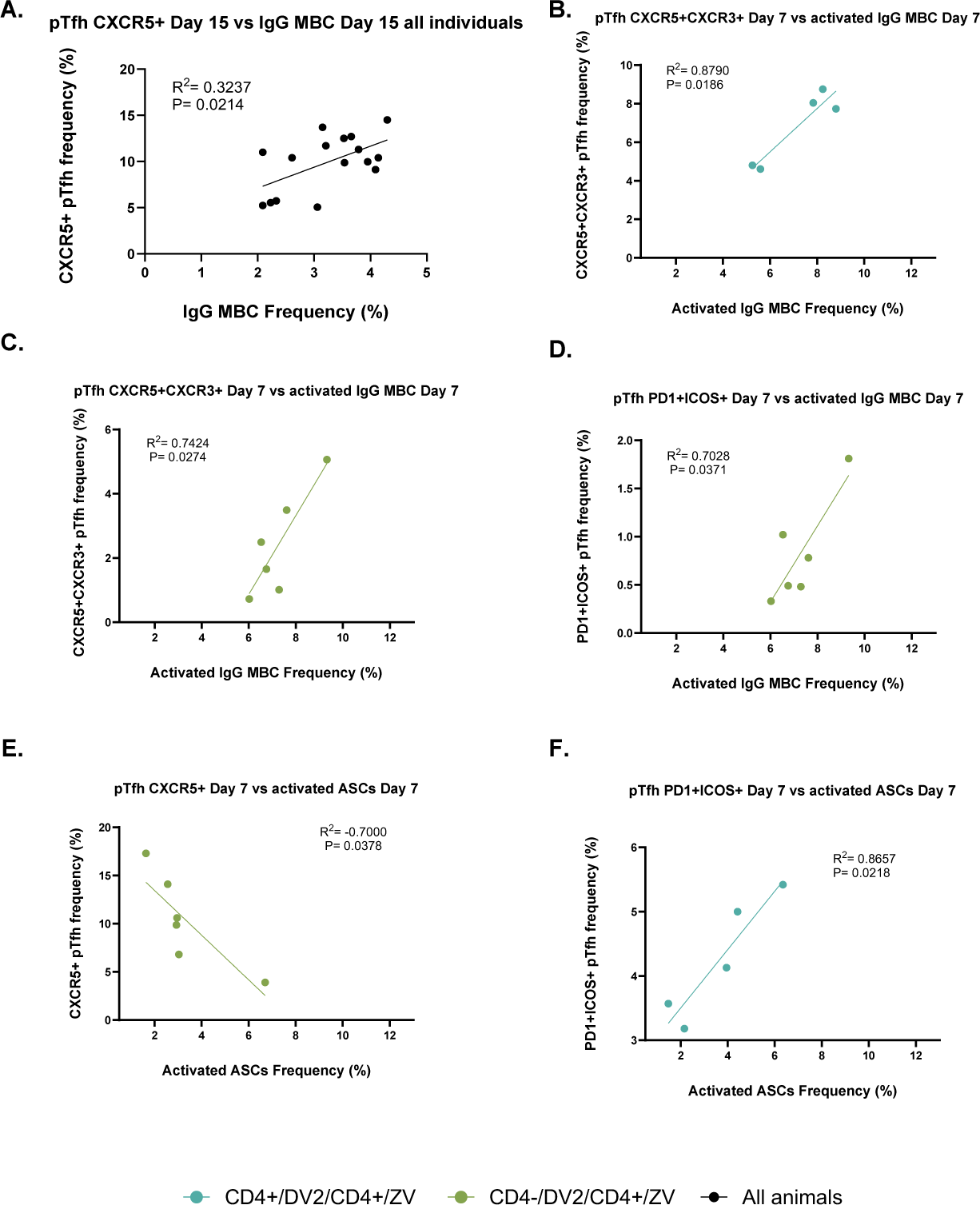
Frequencies of pTfh cells correlate with B cell response cell subsets such as total IgG^+^ MBCs, activated IgG^+^ MBCs, and activated ASCs during the secondary ZIKV infection. Pearson correlations and simple linear regressions were performed on data from pTfh and B cell subsets. All experimental animals are shown in black. Animal cohorts are depicted as follows based on treatment regime: CD4+/DV2/CD4+/ZV (shown in turquoise) and CD4-/DV2/CD4+/ZV (shown in green). **(A)** The levels of total pTfh cells of the CXCR5^+^ phenotype (shown as X5+ in graphs), measured via Flow Cytometry from PBMCs, isolated at day 15 p.c., correlated with frequencies of total IgG^+^ MBCs, measured via Flow Cytometry from isolated whole blood at day 15 p.c.. **(B-C)** Correlations were performed comparing CXCR5^+^CXCR3^+^ pTfh cells (shown as X5+X3+) from day 7 p.c. with **(B)** activated IgG^+^ MBCs from day 7 p.c., in the CD4+/DV2/CD4+/ZV group, and in the **(C)** CD4-/DV2/CD4+/ZV group. Correlations were performed comparing PD1+ICOS+ pTfh cells, from day 7 p.c., with **(D)** activated IgG^+^ MBCs in the CD4-/DV2/CD4+/ZV group. A correlation was performed comparing CXCR5^+^ (X5+) pTfh cells, from day 7 p.c., with **(E)** activated ASCs in the CD4-/DV2/CD4+/ZV group. A correlation was performed comparing PD1+ICOS+ pTfh cells, from day 7 p.c., with **(F)** activated ASCs in the CD4+/DV2/CD4+/ZV group (+/+). Pearson’s rank correlation coefficients were used to calculate correlations, and R^2^- and P-values are indicated. Linear regression is shown as lines and each dot represents one animal.

**SFig 7.**
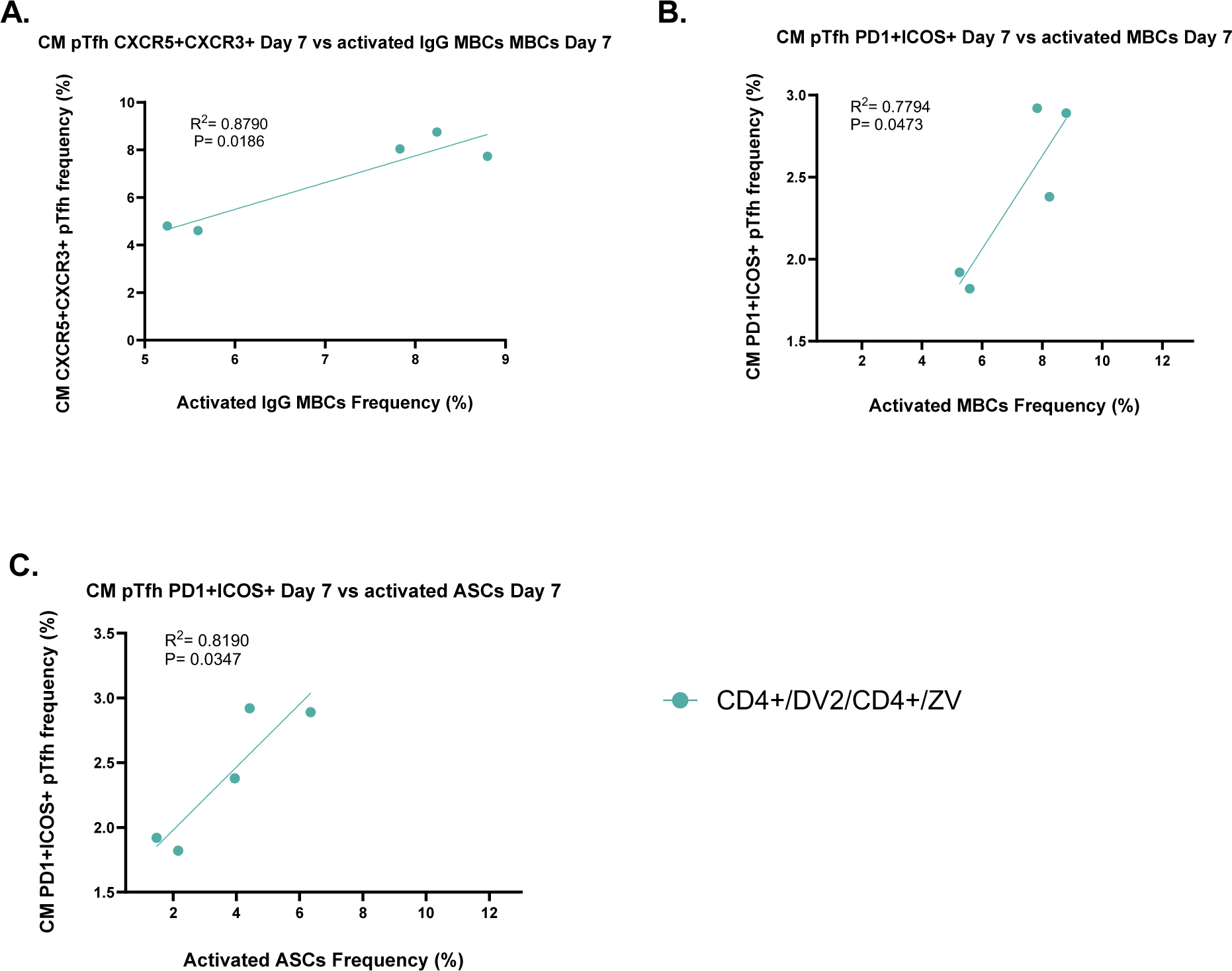
Frequencies of CM pTfh cell subsets correlate with B cell response cell subsets. Pearson correlations and simple linear regressions were performed using data from B cell subsets such as activated IgG^+^ MBCs and ASCs during the secondary ZIKV infection. Animal cohorts are depicted as follows based on treatment regime: CD4+/DV2/CD4+/ZV (shown in turquoise). All significant correlations with CM pTfh cells were only observed in the CD4+/DV2/CD4+/ZV group. Correlations were performed and observed for the CM CXCR5^+^CXCR3^+^ pTfh cells (shown as X5+X3+), from day 7 p.c., with **(A)** activated IgG^+^ MBCs, from day 7 p.c. in the CD4+/DV2/CD4+/ZV group. Correlations were performed for comparing CM PD1+ICOS+ pTfh cells, from day 7 p.c., with **(B)** activated IgG^+^ MBCs in the CD4+/DV2/CD4+/ZV group. Correlations were performed for CM PD1+ICOS+ pTfh cells with **(C)** activated ASCs, from day 7 p.c., in the CD4+/DV2/CD4+/ZV group. Pearson’s rank correlation coefficients were used to calculate correlations, and R^2^- and P-values are indicated. Linear regression is shown as lines and each dot represents one animal.

**SFig 8.**
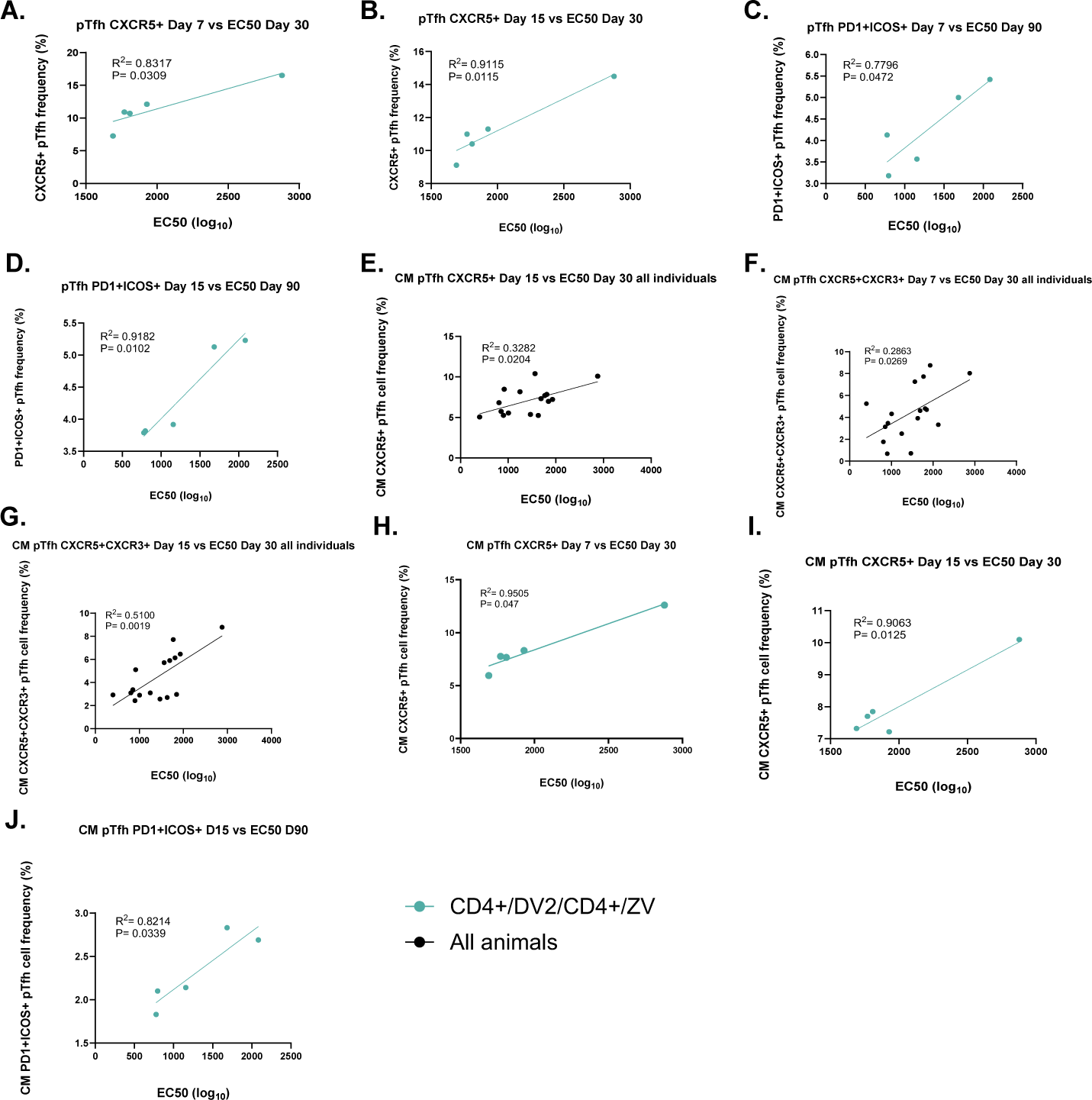
Frequencies of pTfh cell subsets correlate with EC50 (log_10_) values. Data obtained from Focus/Plaque Reduction Neutralization Tests on collected sera from days 30 and 90 p.c. during secondary ZIKV infection was analyzed using Pearson and simple linear regression with pTfh cells. All experimental animals are shown in black. Animal cohorts are depicted as follows based on treatment regime: CD4+/DV2/CD4+/ZV (shown in turquoise). **(A-B)** Correlations were performed comparing CXCR5^+^ pTfh cells (shown as X5+), from **(A)** days 7 and **(B)** days 15 p.c. with EC50 values from day 30 p.c., in the CD4+/DV2/CD4+/ZV group. **(C-D)** Correlations were performed comparing PD1+ICOS+ pTfh cells from **(C)** day 7 and **(D)** day 15 p.c. with EC50 values from day 90 p.c., in the CD4+/DV2/CD4+/ZV group. **(E-I)** Frequencies of CM pTfh cell subsets correlate with EC50 values from days 30 and 90 p.c.. **(E-G)** Correlations were performed using all experimental comparing **(E)** CM CXCR5^+^ (X5+) pTfh cells, from day 15 p.c., with EC50 values from day 30 p.c., comparing **(F)** CM CXCR5^+^CXCR3^+^ (shown as X5+X3+) pTfh cells, from day 7 p.c., with EC50 values from day 30 p.c., and comparing **(G)** CM CXCR5^+^CXCR3^+^ (X5+X3+) pTfh cells, from day 15 p.c.. Correlations were performed comparing CM CXCR5^+^ (X5+) pTfh cells, from **(H)** day 7 and **(I)** day 15 p.c., with EC50 values from day 30 p.c.. Correlations were performed comparing CM PD1+ICOS+ pTfh cells, from day 15 p.c., with **(J)** EC50 values from day 90 p.c.. Pearson’s rank correlation coefficients were used to calculate correlations, and R^2^- and P-values are indicated. Linear regression is shown as lines and each dot represents one animal.

**SFig 9.**
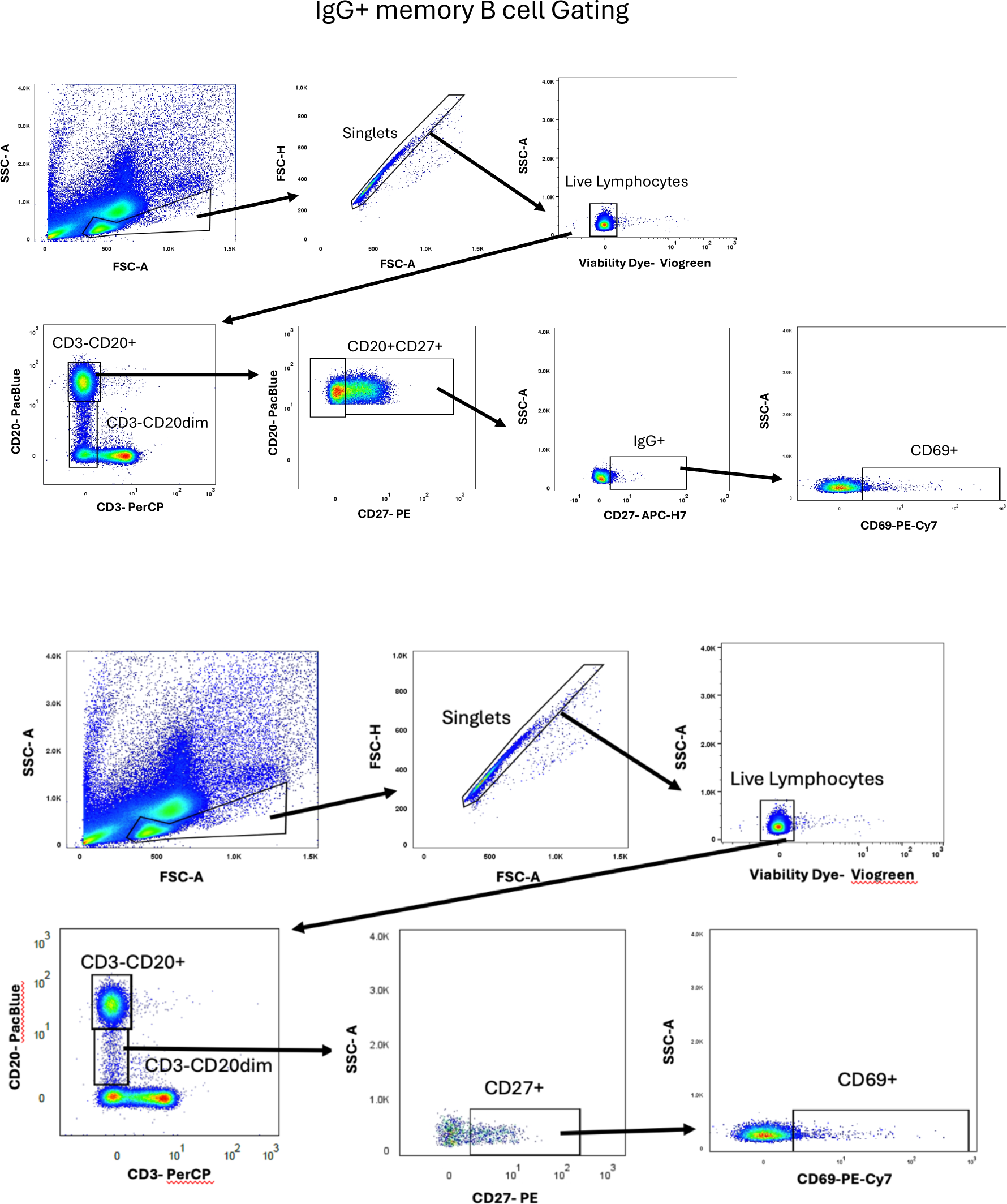
Gating strategy for B cells. Gating strategy used to define B cells and subsets. Lymphocytes were gated based on their characteristic forward and side scatter pattern (FSC, SSC). B cells were defined as CD20+CD3-.

**SFig 10.**
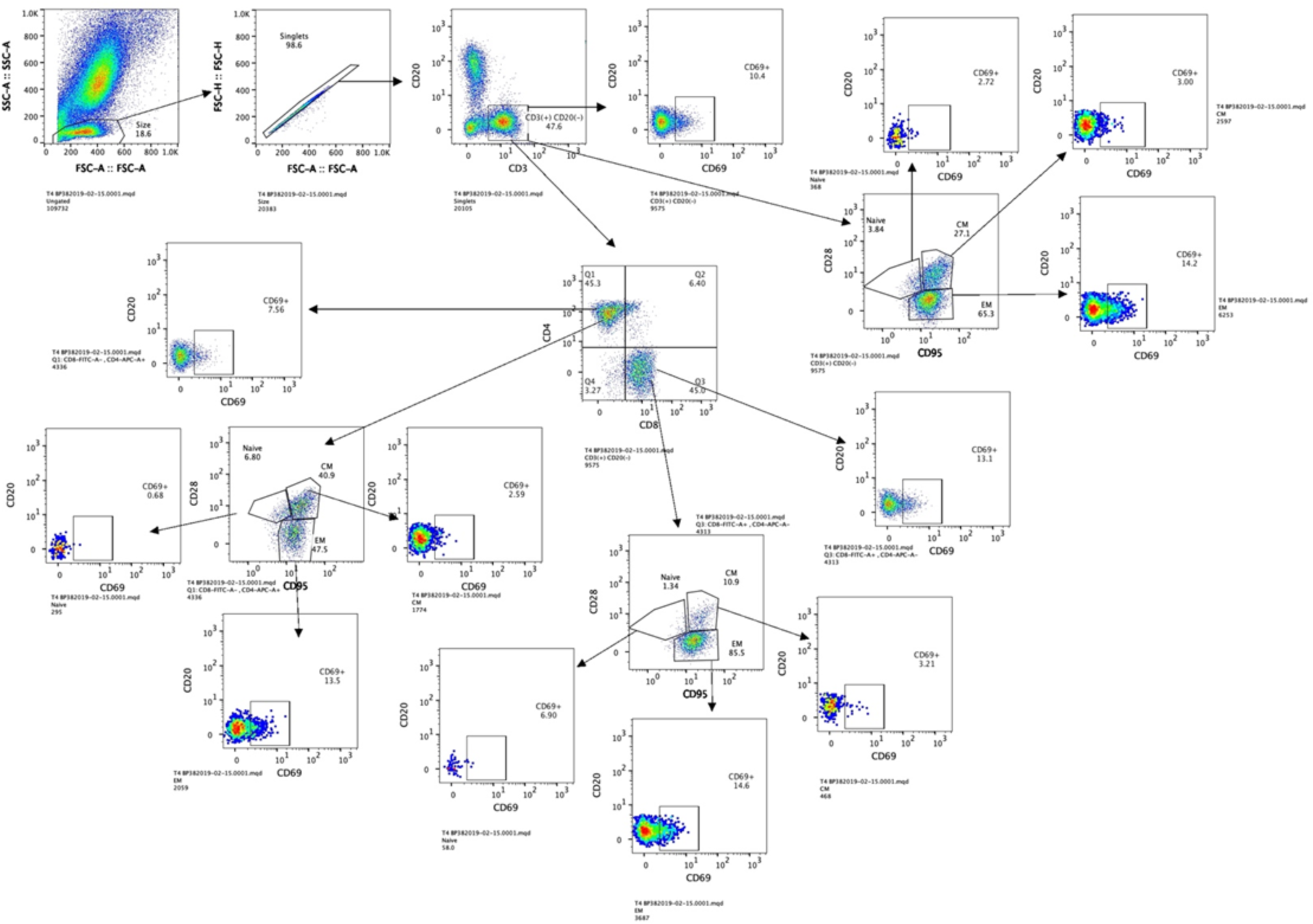
Gating strategy for T cells. Gating strategy used to define T cells and subsets. Lymphocytes were gated based on their characteristic forward and side scatter pattern (FSC, SSC). T cells were defined as CD3+CD20+. CD4+ and CD8+ T cells were defined as CD3+CD4+ and CD3+CD8+, respectively.

**SFig 11.**
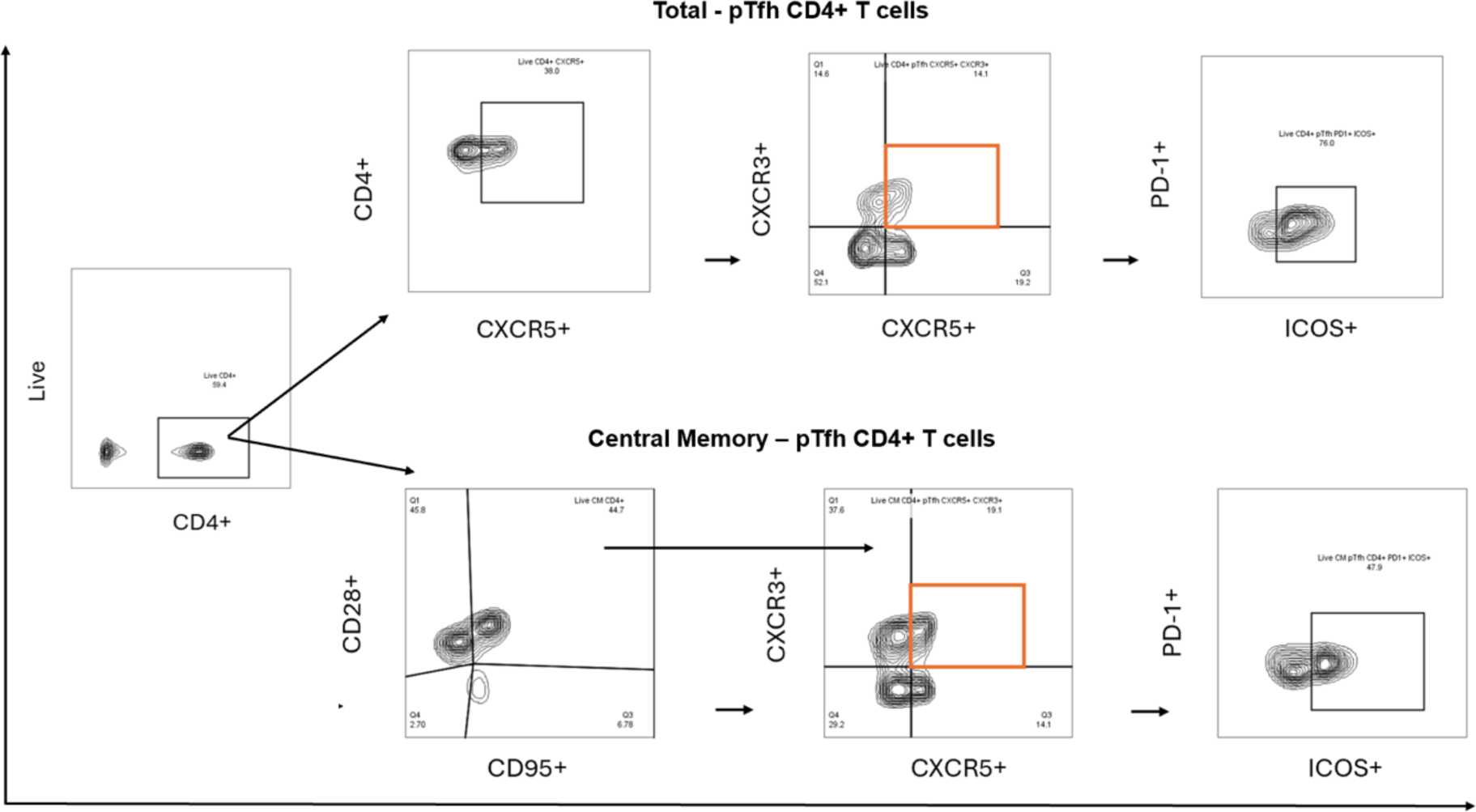
Gating strategy for pTfh CD4+ T cells. Gating strategy used to define pTfh CD4+ T and total central memory (CM-pTfh) CD4+ T cells based on CD28+, CD95+, CXCR5+, CXCR3+, PD1+, and ICOS+ surface markers expression.

**Table S3.** Parameters used to perform PCA analysis. PCA analysis was performed using parameters from days 7, 15, 30 and 90 post ZIKV infection.

| **Parameter** | **Day of measurement for PCA** |
| --- | --- |
| TFH_CD4_ICOS_PD1 | Day 7, Day 15 |
| zikav_od_igm | Day 15, Day 30 |
| zikav_od_igg | Day 15, Day 30, Day 90 |
| denv_od_igm | Day 15 |
| denv_od_igg | Day 15, Day, 30, Day 90 |
| neutralization vs. ZikaV | Day 15, Day 30, Day 90 |
| neutralization vs. DengueV | Day 15, Day 30, Day 90 |
| CD4_T_cells | Day 15, Day 30, Day 90 |
| CD8_T_cells | Day 15, Day 30, Day 90 |
| T4_CM | Day 15, Day 30, Day 90 |
| T4_CM_CD69 | Day 15, Day 30, Day 90 |
| T8_CM | Day 15, Day 30, Day 90 |
| T8_CM_CD69 | Day 15, Day 30, Day 90 |
| T4_EM | Day 15, Day 30, Day 90 |
| T4_EM_CD69 | Day 15, Day 30, Day 90 |
| T8_EM | Day 15, Day 30, Day 90 |
| T8_EM_CD69 | Day 15, Day 30, Day 90 |
| CD4_Naive | Day 15, Day 30, Day 90 |
| CD4_Naive_CD69 | Day 15, Day 30, Day 90 |
| CD8_Naive | Day 15, Day 30, Day 90 |
| CD8_Naive_CD69 | Day 15, Day 30, Day 90 |
| B_Cells | Day 15, Day 30, Day 90 |
| B_Cells_CD69 | Day 15, Day 30, Day 90 |
| CD27_CD69 | Day 15, Day 30, Day 90 |
| CD20_CD27 | Day 15, Day 30, Day 90 |
| EDIII_CRI_D1 | DAY 90 |
| EDIII_CRI_D2 | DAY 90 |
| EDIII_CRI_D3 | DAY 90 |
| EDIII_CRI_D4 | DAY 90 |
| EDIII_CRI_Z | DAY 90 |
| NS1_CRI_D1 | DAY 90 |
| NS1_CRI_D2 | DAY 90 |
| NS1_CRI_D3 | DAY 90 |
| NS1_CRI_D4 | DAY 90 |
| NS1_CRI_Z | DAY 90 |
| OD = Optical Density |  |
| CM = Central Memory |  |
| tfh = T Follicular Helper |  |
| CRI = Cross Reactivity Index |  |

